# Multilevel engineering of cyanobacterial energy metabolism advances photosynthetic hydrogen production while revealing its constraints

**DOI:** 10.64898/2026.07.21.739770

**Authors:** Marvin Amadeus Itzenhäuser, Samuel Grimm, Lara Ruprecht, Johanna Wiedener, Fabian Brandenburg, Ron Stauder, Pål William Wallace, Jens Olaf Krömer, Andreas Schmid, Stephan Klähn

## Abstract

Hydrogen (H_2_) is a promising sustainable energy carrier, and its direct production from photosynthetic water splitting is appealing. Yet long-term photosynthetic hydrogen production by cyanobacteria remains inefficient despite decades of engineering. Here, we systematically dissect the hierarchical and state-dependent constraint architecture governing sustained H_2_ evolution in *Synechocystis* sp. PCC 6803. We show that hydrogenase overexpression relieves the primary enzymatic limitation, exposing ATP/NADPH balancing and competing electron sinks as successive metabolic constraints. Inspired by cyanophage strategies, we engineered synthetic CP12-based regulatory proteins that redirect photosynthetic electrons from CO_2_ fixation toward H_2_ production. Combining these interventions increases H_2_ production by over two orders of magnitude relative to the previous benchmark system, demonstrating that sustained H_2_ production requires coordinated management of metabolism and regulation rather than elimination of a single bottleneck. However, overcoming these constraints also promotes metabolic adaptations and genetic instability, illustrating the trade-off between maximal H_2_ production and long-term metabolic stability.

## Introduction

Cyanobacteria perform oxygenic photosynthesis, coupling solar energy capture with water oxidation and an autotrophic metabolism based on CO_2_ fixation. The ability to directly channel solar power into product formation positions cyanobacteria as promising chassis for sustainable biotechnology, and research in this area has intensified considerably in recent years ^1^. In contrast to conventional chemoheterotrophic production systems, photobiotechnological approaches based on cyanobacteria bypass the need for plant-derived carbohydrates. Thereby reliance on arable land could be reduced, food-versus-fuel concerns alleviated, and favourable light-to-biomass conversion efficiencies compared to terrestrial plants can be employed ^2–6^. The metabolic diversity and genetic accessibility of cyanobacteria have enabled production of diverse compounds, including fine chemicals ^2,4,7^, bulk products ^8–10^, and energy carriers such as ethanol or hydrogen (H_2_) ^11–13^. More recently, engineered cyanobacteria have also been explored as light-driven whole-cell biocatalysts for redox-intensive transformations powered exclusively by photosynthetic electrons ^14–16^.

Despite these advances, the overall productivities with respect to the H_2_-formation rate and its stability remained limited. A central challenge lies in the intrinsic prioritization of growth and metabolic homeostasis ^17,18^. This competes with product formation, often leading to low energy efficiencies and suboptimal carbon partitioning ^19^. Photosynthetically generated ATP and NADPH are tightly allocated to sustain biomass formation and redox balance ^20^. Thus, redirecting carbon and reducing power toward product formation requires overcoming the intrinsic prioritization of growth and redox stability ^1^. Metabolic engineering strategies have therefore focused on introducing heterologous pathways, enhancing enzyme abundance, or eliminating competing reactions ^21^. While such interventions can increase product yields, they often rely on static genetic modifications and do not fully account for the dynamic regulatory architecture governing photosynthetic metabolism.

Microbial metabolism operates as a highly responsive and tightly regulated network that continuously adapts to environmental and intracellular cues. Beyond transcriptional and translational regulation of genes, enzyme activities are modulated by kinetics, metabolites, covalent modifications, or interaction with regulatory proteins, enabling rapid adjustment of metabolic fluxes ^22^. Deliberate alteration of gene expression has become a fundamental strategy in metabolic engineering, that, however, only changes the abundance of specific proteins and not necessarily global metabolic fluxes. Targeting intrinsic regulatory mechanisms may therefore offer a powerful yet largely underexplored strategy to dynamically reprogram metabolism. In this context, small proteins of fewer than 100 amino acids have emerged as potent regulators of metabolic processes in bacteria that can be used to alter broad regulatory mechanisms with low metabolic burden for their production ^23,24^. In cyanobacteria, small regulatory proteins modulate central pathways including nitrogen assimilation and carbon partitioning ^25–27^. For example, regulatory control of carbon flux through the small regulatory protein PirC (also designated as CfrA) can substantially alter metabolite routing and enable marked increases in accumulation of polyhydroxybutyrate or sucrose ^28–30^. These findings highlight the capacity of small regulatory proteins to exert disproportionate control over metabolic flux distribution ^31^. Despite their demonstrated regulatory potency, the rational redesign of small proteins as synthetic tools for dynamic metabolic engineering remains largely untapped.

In cyanobacteria, CO_2_ is fixed via the Calvin-Benson-Bassham (CBB) cycle. Its activity is dynamically controlled by the small, redox-responsive protein CP12, which forms reversible complexes with glyceraldehyde-3-phosphate dehydrogenase (GAPDH) and phosphoribulokinase (PRK), thereby modulating CO_2_ fixation in response to the cellular redox state ^32–34^. Given the high energetic demand of CO_2_ fixation, CP12 effectively governs a major competing sink for photosynthetic reducing power ^18,35^. Previous studies have demonstrated that deletion of *cp12* can enhance the production of carbon-based compounds such as 2,3-butanediol or terpenoids, underscoring its potential as a regulatory engineering target ^36,37^. Among photobiotechnological applications, H_2_ formation constitutes a special case. Unlike carbon-based products, H_2_ formation depends exclusively on the availability of protons and reducing equivalents and hence, does not require carbon flux through the CBB cycle. From this perspective, CP12 represents a particularly attractive regulatory handle for H_2_ production as dynamic inhibitor of the CBB cycle.

The conceptual appeal of directly coupling photosynthetic water oxidation to proton reduction via hydrogenases has motivated extensive engineering efforts over the past decades. Strategies have included modification of electron transport pathways ^38^, overexpression of the native ^39^ and introduction of heterologous hydrogenases ^40,41^, elimination of competing sinks ^42^, a hydrogenase - photosystem I fusion to channel electrons directly from the light reaction into H_2_ formation ^11^, or encapsulation of hydrogenases in carboxysomes for protection from oxygen ^43^. Despite these extensive efforts, H_2_ formation rates remain far below theoretical expectations and are often not sustained over extended periods. Importantly, most strategies have been developed and evaluated in isolation, and systematic integration of impacting parameters has been limited. It therefore remains unclear whether the modest gains of performance reflect incomplete combinatorial optimization, unresolved limitations, or emergent properties of the photosynthetic network that actively preserve redox homeostasis and restrict sustained large-scale diversion of electron flux.

Here, we pursued a system-level strategy to reallocate photosynthetic electron flux toward sustained H_2_ production in the cyanobacterium *Synechocystis* sp. PCC 6803 (hereafter *Synechocystis*). We employ an oxygen removal approach enabling long-term activity of the intrinsic, bidirectional, oxygen-sensitive [NiFe]-hydrogenase that enables H_2_ evolution with electrons derived from water. To interrogate the coupling between ATP and reductant generation on H_2_ production, we further evaluated the impact of modulating the proton motive force via an uncoupler. Most importantly, we designed synthetic variants of the small regulatory protein CP12 to inhibit the CBB cycle under light conditions, thereby targeting the dominant endogenous sink for photosynthetic reducing power. Integration of these regulatory and metabolic interventions with modifications of the photosynthetic electron transport chain and enhanced hydrogenase activity established a robust, sustained hydrogen-producing regime productive over days, reaching an average rate of 0.61 µmol min^-1^ L^-1^ OD^-1^. However, driving the photosynthetic system and central metabolism toward elevated H_2_ production uncovered pronounced fitness trade-offs and recurrent genetic instability, exposing the inherent resistance of photosynthetic metabolism to sustained redirection of its core fluxes.

## Results

### Sustained H_2_ production with electrons from water is achieved in a bacterial co-culture system ensuring anaerobicity

The wild type (WT) of *Synechocystis* produces H_2_ via its bidirectional [NiFe]-hydrogenase either fermentatively in the dark, in particular in the presence of external, metabolizable carbon sources or transiently during the transition from darkness to light ^38^. This transient production phase is attributed to the rapid reduction of cellular redox pools upon illumination while CO_2_ fixation via the CBB cycle, the major electron sink **(Fig. 1A)**, remains temporarily inactive due to inhibition by the redox-sensitive protein CP12 ^44^. During this lag phase, NADP and ferredoxin pools become sufficiently reduced to reach redox potentials permissive for transient, light-driven H_2_ formation **(Fig. 1B)**. Upon activation of the CBB cycle electron flux is redirected toward CO_2_ fixation, resulting in a steady state in which the intracellular redox potential favors H_2_ consumption rather than production, which is why H_2_ levels subsequently decrease ^45^.

**Figure 1:**
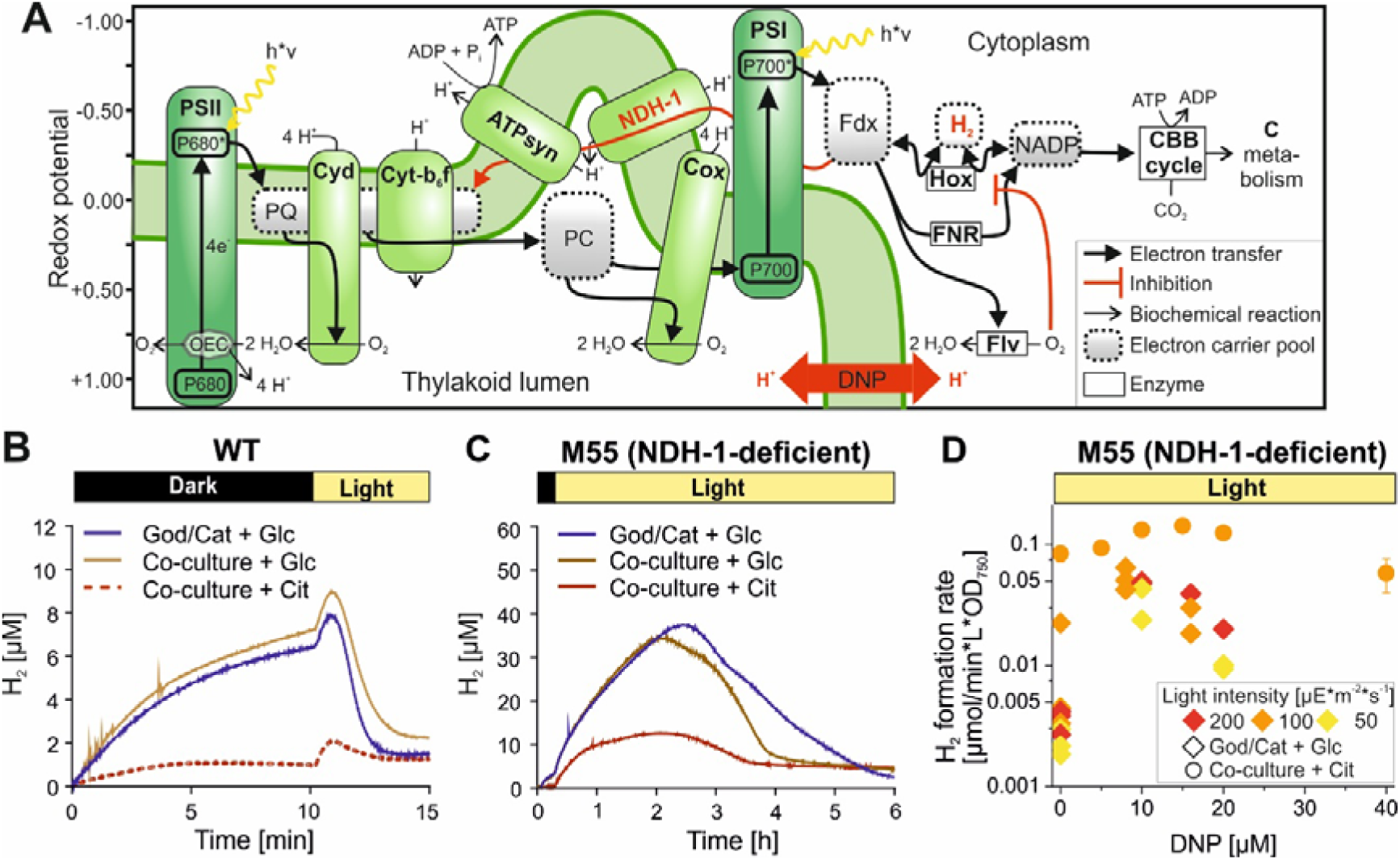
Light-dependent H_2_ production by *Synechocystis* in different experimental and genetic configurations. **A**: Schematic of the photosynthetic electron transport chain and CO_2_ fixation in cyanobacteria. Heights in the schema represent redox potentials. The redox potential of electron carrier pools depends on the ratio of its reduced to its oxidized form and are therefore depicted as areas ranging from 99.99% reduced to 99.99% oxidized. PSI/PSII: photosystem I or II; PQ: plastochinone; Cyd: cytochrome *bd* quinol oxidase, Cyt-b_6_f: cytochrome b_6_f complex; ATPsyn: ATP synthase; PC: plastocyanin; NDH-1: NAD(P)H dehydrogenase-like complex type 1; Cox: Cytochrome c oxidase; Fdx: ferredoxin; Flv: flavodiiron proteins; DNP: 2,4-dinitrophenol; Hox: [NiFe]-hydrogenase; FNR: ferredoxin-NADP^+^ oxidoreductase. **B,C:** H_2_ evolution during light and dark phases in *Synechocystis* WT and the M55 mutant in which the *ndhB* gene is disrupted. The measurements were conducted *in vivo* using UniSense electrodes. To maintain anaerobic conditions also under photosynthetic conditions (light phase) cultures were supplemented with 10 mM glucose or citrate and either an enzyme mixture of 40 U glucose oxidase and 50 U catalase (God/Cat) or a defined amount of *Pseudomonas taiwanensis* VLB120 cells. A hydrogenase-negative *Synechocystis* strain (Δ*hox*) was tested in the same setup and did not evolve any hydrogen proving that it is not coming from *Pseudomonas* cells (not shown). Please note the different time resolution and H_2_ yields in B and C. The obtained yields enabled H_2_ quantification via gas chromatography. **D:** H_2_ formation rates of the *Synechocystis* M55 in co-culture with *Pseudomonas* and 10 mM citrate or 10 mM glucose and God/Cat (40/50 U) quantified by gas chromatography after ∼ 48 h or ∼24 h in light, respectively. DNP was added in different concentrations to dissipate the proton gradient over the thylakoid membrane. Data represent individual measurements (God/Cat) or the mean ± SD of three biological replicates (Co-culture); the entire dataset is given in Supplementary Table S1.

Most hydrogenases, including the [NiFe]-hydrogenase of *Synechocystis,* are oxygen sensitive ^46^. To enable hydrogenase activity under photosynthetic conditions, oxygen that is produced by photosystem II (PSII) must be continuously removed. Previous studies employed an enzymatic oxygen scavenging system consisting of glucose oxidase (God) and catalase (Cat), which oxidizes glucose added to the medium ^11,38,45^. However, the supplemented glucose can be metabolized by *Synechocystis,* thereby inducing a mixotrophic state that enables H_2_ production in darkness (**Fig. 1B**). In the light, photosynthesis additionally provides reducing equivalents. The [NiFe]-hydrogenase of *Synechocystis* accepts electrons from ferredoxin and NAD(P)H ^38,47^. Because NAD(P)H can be generated directly from glucose oxidation and fed into the photosynthetic electron transport chain via the NAD(P)H dehydrogenase-like complexes (NDH, **Fig. 1A**) this assay does not allow discrimination between electrons derived from glucose catabolism and those originating from water splitting at PSII.

To avoid supplying *Synechocystis* with a metabolizable organic carbon source, we additionally used an assay that we describe in detail in companion publications^48,49^ and provide here only essential features relevant for this study. The assay relies on respiratory oxygen consumption by *Pseudomonas taiwanensis* VLB120 (hereafter *Pseudomonas*) in a microbial consortium with *Synechocystis*. Co-cultures supplied with glucose as carbon-source for *Pseudomonas* exhibited H_2_ production kinetics comparable to the God/Cat enzymatic assay **(Fig. 1B)**, confirming effective oxygen removal. When citrate was supplied as the sole organic carbon source, which is metabolized by *Pseudomonas* but not by *Synechocystis* ^48,50^, fermentative H_2_ production in the dark was largely abolished, while the transient production phase immediately after illumination and the subsequent re-uptake remained detectable **(Fig. 1B)**.

In addition, to achieve prolonged production, we employed the NDH-1-deficient *Synechocystis* mutant “M55” (hereafter referred to as M55), previously reported to exhibit extended light-driven H_2_ evolution compared to WT ^38^. When tested under the different oxygen removal strategies, the M55 mutant produced substantially higher amounts of H_2_ compared to the WT over several hours (**Fig. 1B,C**). Although glucose supplementation – either in the God/Cat enzymatic assay or in glucose-fed co-cultures – resulted in higher dissolved H_2_ concentrations in the short term, the citrate-based co-culture exhibited more sustained H_2_ production in the light and markedly reduced fermentative H_2_ production in the dark (**Fig. 1C**). This is in-line with the finding that electrons from glucose oxidation by *Synechocystis* significantly contributes to both light-driven and fermentative H_2_ production in the M55 mutant and electrons for H_2_ formation originate from water-splitting at PSII only if citrate is used in the co-culture assay ^48^. Interestingly, although electrode measurements indicated lower dissolved H_2_ levels in citrate-fed co-cultures, headspace analysis after extended periods (> 6h) revealed substantially higher H_2_ production rates compared to the glucose-based enzymatic assay **(Fig. 1D)**. Together, these results establish the citrate-fed co-culture system described in the companion studies ^48,49^ as a suitable platform for analysing sustained photosynthetic H_2_ formation by *Synechocystis* under anaerobic conditions without providing the cyanobacterium with a metabolizable organic carbon source.

### Using a chemical uncoupler to modulate the proton motif force towards enhanced H_2_ production

In the M55 mutant the *ndhB* gene is disrupted, preventing assembly of a functional NDH-1 complex ^38^. The NDH-1 complex normally transfers electrons from ferredoxin or NADPH to the plastoquinone pool while concomitantly translocating protons from the cytoplasm into the thylakoid lumen (**Fig. 1A**). Through this reaction, cyclic electron flow around PSI contributes to the generation of proton motive force (pmf), thereby dissipating excess reducing equivalents, supporting ATP synthesis and balancing the ATP/NADPH ratio required for CO_2_ fixation and other anabolic reactions. The NDH-1 deletion changes this balance and ATP production becomes limiting relative to NADPH generation and redox dissipation is prevented ^38,51,52^. Under anaerobic conditions, dissipation of excess reducing equivalents through oxygen reduction is likewise abolished. As a consequence, the NADPH pool becomes highly reduced, a condition previously linked to the sustained H_2_ formation phenotype of the M55 mutant ^38^. Thus, in anaerobic cultures of the M55 mutant, H_2_ formation can be employed to partially restore the ATP/NADPH balance by functioning as an alternative electron sink. Importantly, the linkage between H_2_ formation and the ATP/NADPH balance could also constrain H_2_ formation rates. If H_2_ evolution exceeds the rates necessary to restore the ATP/NADPH balance, pmf buildup would lead to feedback inhibition reducing photosynthetic activity ^53^.

As H_2_ production is linked to the ATP/NADPH balance, we modulated the pmf using the protonophore 2,4 dinitrophenol (DNP), which facilitates proton translocation across biological membranes along the concentration gradient ^54^. Different concentrations of DNP were applied to M55 cultures in both the glucose-based enzymatic oxygen scavenging assay and the citrate-fed co-culture system (**Fig. 1D**). In both assays, DNP exerted a concentration-dependent stimulatory effect on H_2_ production. In the enzymatic assay with glucose, 8 µM DNP increased H_2_ production rate by approximately tenfold achieving almost the same rates as in the citrate-fed co-culture system. Also, in the citrate-fed co-culture setup, 15 µM DNP further enhanced H_2_ production rates by a factor of 1.7 **(Fig. 1D)**. Notably, in this system the stimulatory effect became pronounced only after extended incubation (approx. 48 hours). Collectively, these results demonstrate that the pmf represents a bioenergetic constraint on sustained H_2_ production and that partial uncoupling can substantially enhance H_2_ evolution dependent on the prevailing metabolic state of the cells.

### Overproduction of the intrinsic [NiFe]-hydrogenase increases H_2_ production capacity in *Synechocystis*

Achieving high H_2_ production rates requires sufficient catalytic capacity of the intrinsic [NiFe]-hydrogenase encoded by the genes *hoxEFUYH*. Transcription of this operon is controlled by the regulator LexA ^55^, for which, however the signalling cascade remains largely unresolved in *Synechocystis*. Notably, in our hands H_2_ production rates varied considerably (relative standard deviation of 118 %) between biological replicates under otherwise identical conditions which might be caused by variable expression of *hoxEFUYH.* Insufficient expression may also limit H_2_ production. Previous studies demonstrated that overexpression of *hoxEFUYH* in a WT background increased the [NiFe] hydrogenase activity up to threefold ^39^. However, extended H_2_ production in the WT remained prevented by the high steady-state redox potential. In contrast, whether catalytic capacity limits H_2_ evolution in the overreduced M55 background has not been tested. The M55 mutant is based on a *Synechocystis* substrain different to the WT used in this study, complicating direct comparisons as genetic and phenotypic variances have been noticed in *Synechocystis* lab substrains ^56^. To address this, we used a Δ*ndhB* strain in which the disrupted *ndhB* gene and the antibiotic resistance cassette from the M55 mutant was amplified and used this to establish the same *ndhB* disruption in the here employed WT background ^48^. Subsequently, we deleted the entire *hox* operon to obtain the Δ*ndhB* Δ*hox* strain **(Fig. 2A)**. Full Segregation of the respective strains was verified using PCR **(Fig. S1)**. The coding sequences of the *hox* operon were then reintroduced on replicative plasmids into that strain under the control of three constitutive promoters of different strength producing the Δ*ndhB* Δ*hox* + P_J231XX_*::hoxEFUYH* strains **(Fig. 2A)**.

**Figure 2:**
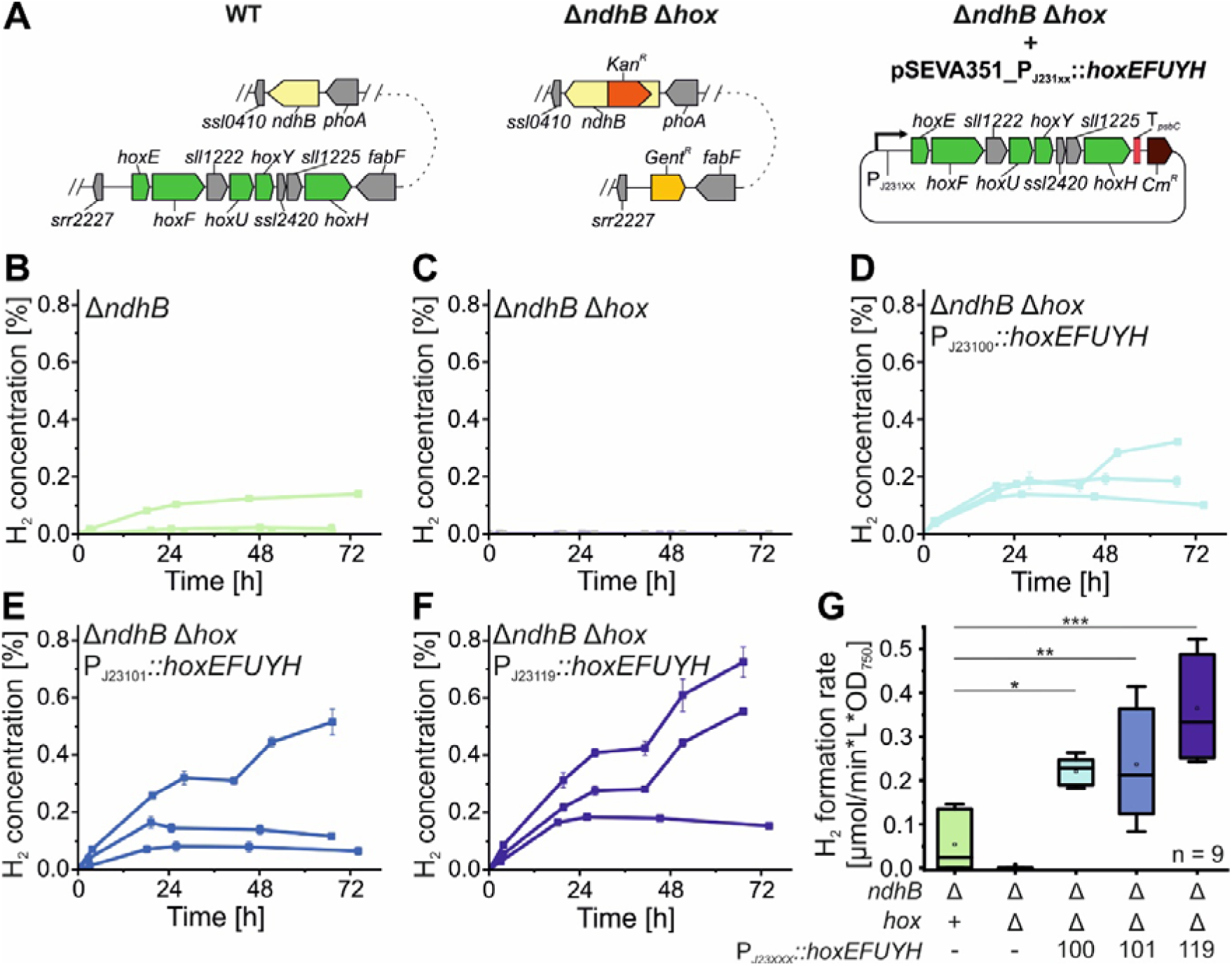
Hydrogen production of cyclic electron transport deficient *Synechocystis* strains overexpressing the hydrogenase genes. **A**: Genetic setup of the presented strains. **B-F:** H_2_ concentration in the headspace of sealed citrate-fed co-cultures with *Pseudomonas* over three days of incubation in light. Separate lines represent independent experiments with data points indicating averages of measurements of three biological replicates ± SD**. G:** H_2_ production rates in the initial 24 h normalized to culture density and volume. The box plots show median (line), average (square), 25 – 75% quartiles (boxes), and minimum and maximum values within 1.5× the interquartile range (whiskers). Statistical significance between groups was assessed using a Tukey post hoc test and is indicated as follows: p□<□0.05 (*), p□<□0.01 (**), p□<□0.001 (***).

The Δ*ndhB* strain exhibited H_2_ production comparable to the original M55 mutant, whereas the Δ*ndhB* Δ*hox* strain showed no detectable H_2_ formation **(Fig. 2B,C)**, confirming that H_2_ evolution is strictly dependent on the intrinsic hydrogenase. Reintroduction and overexpression of the *hoxEFUYH* operon increased H_2_ production up to sevenfold relative to the Δ*ndhB* parental strain. Thereby, H_2_ formation positively correlated with promoter strength **(Fig. 2D-G)** as the highest rates were achieved when *hoxEFUYH* expression was driven by PJ23119 which was previously proven to mediate stronger expression in *Synechocystis* compared to P23101 and PJ23100 ^57^. This indicates that catalytic capacity represents a quantitative determinant of H_2_ evolution under these conditions. Interestingly, while in the Δ*ndhB* strain the H_2_ formation had a variation of 118% across independent experiments **(Fig. 2B)**, it varied only between 16% and 54% for the Δ*ndhB* Δ*hox* P_J231XX_*::hoxEFUYH* strains. However, we also noticed clones of the Δ*ndhB* Δ*hox* P_J231XX_*::hoxEFUYH* strains that nearly lost their H_2_ production ability completely after prolonged cultivation despite retaining the plasmid. This suggests selection pressure against high and uncontrolled expression of the hydrogenase operon (see discussion about genetic instability).

### Utilization of the central CBB cycle regulator CP12 to reduce competition for electrons during photosynthetic H_2_ production

In the WT the delayed activation of the CBB cycle following illumination results in transient over-reduction of the NADPH and ferredoxin pools, which underlies the short burst of H_2_ production observed in a dark-light transition ^58^. This physiological phenomenon indicates that limiting the dominant electron sink of photosynthesis can redirect reducing equivalents toward H_2_ formation. However, permanent attenuation of the CBB cycle would severely compromise cellular fitness. Thus, rather than static suppression, a tool is required that enables rapid and inducible restriction of CBB cycle activity in the light, creating a redox imbalance that can be dissipated via H_2_ production. The small regulatory protein CP12 governs CBB cycle activity and therefore represents a strategic engineering target for this purpose. However, CP12 itself is redox-regulated ^34^. Under oxidizing conditions, two intramolecular disulfide bridges form, stabilizing CP12 in an ordered conformation that enables binding to glyceraldehyde 3-phosphate dehydrogenase (GAPDH) and phosphoribulokinase (PRK), thereby inhibiting their activity ^32^ **(Fig. 3A)**. Upon illumination, reduction via thioredoxin disrupts these disulfide bonds, rendering CP12 unstructured and releasing GAPDH and PRK to enable CO_2_ fixation ^59,60^ **(Fig. 3B)**. Through this mechanism, the cells deactivate the CO_2_ fixation in the dark to prevent a futile cycle of CO_2_ fixation with redox and energy equivalents from the degradation of storage components and dynamically adjust CBB cycle activity in response to the cellular redox state ^44^.

**Figure 3:**
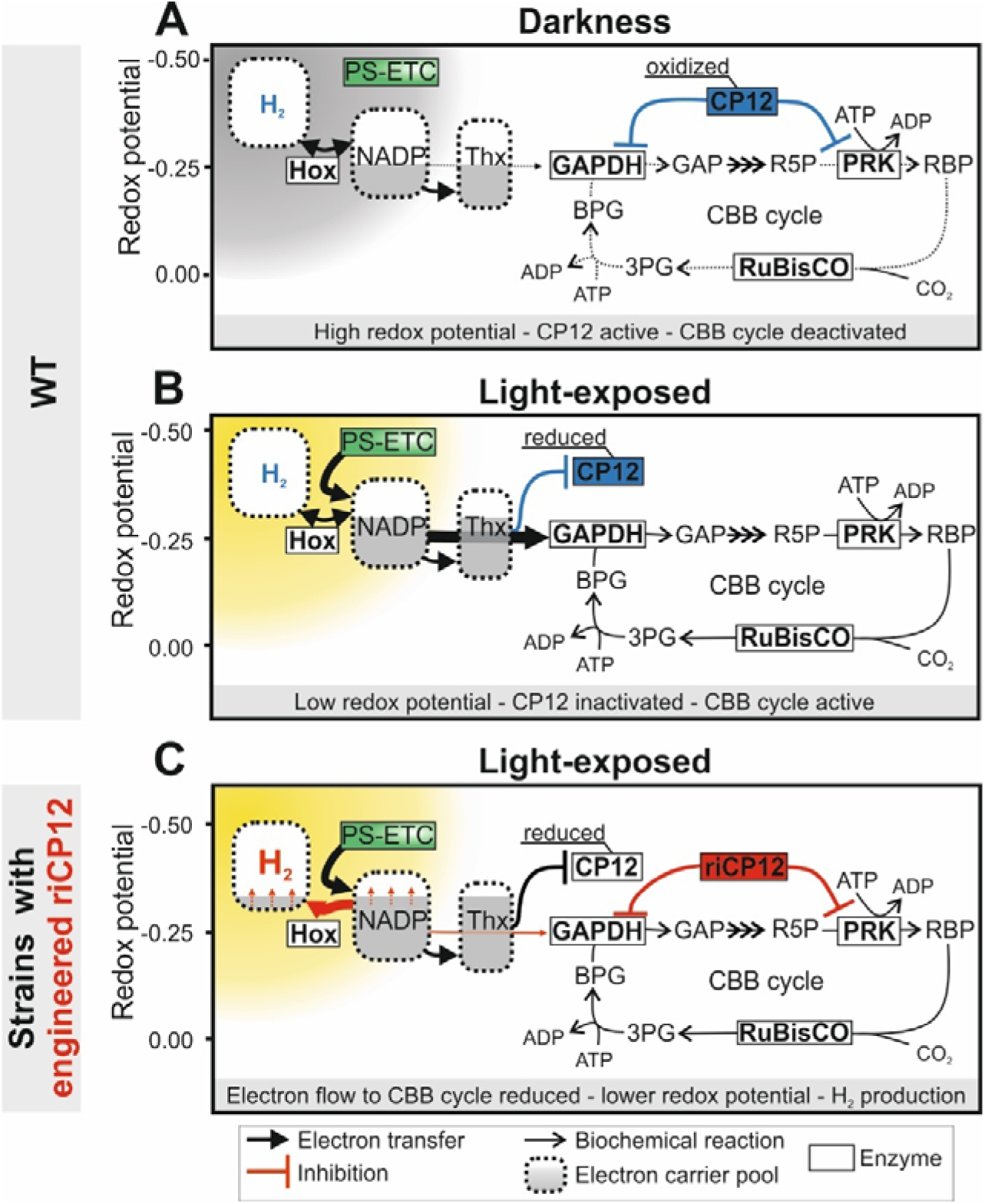
CP12-dependent regulation of the Calvin-Benson-Bassham cycle and its utilization to direct electron flow into H_2_. **A-B**: Schematic of the redox-dependent regulation of the CBB cycle by the small regulatory protein CP12 in dark (A) and light (B) conditions. **C:** Desired effect of the introduction of a redox-insensitive CP12 variant (riCP12) on the photosynthetic H_2_ production. The redox potential of electron carrier pools depends on the ratio of its reduced to its oxidized form and are therefore depicted as areas ranging from 99.99% reduced to 99.99% oxidized. PS-ETC: Photosynthetic electron transport chain; Hox: [NiFe]-hydrogenase; Thx: thioredoxin; GAPDH: glyceraldehyde 3-phosphate dehydrogenase, BPG: 1,3-bisphosphoglycerate; GAP: glyceraldehyde 3-phosphate; 3PG: 3-phosphoglycerate; CP12: chloroplast protein of 12 kDa; RuBisCO: ribulose-1,5-bisphosphate carboxylase/oxygenase; R5P: ribulose 5-phosphate; PRK: phosphoribulokinase; RBP: ribulose-1,5-bisphosphate

Because the endogenous CP12 of *Synechocystis* (CP12^6803^) becomes inactive under reducing conditions, i.e. in the light, it cannot be directly employed to restrict CBB cycle activity during photosynthetic H_2_ production. To establish a regulatory tool capable of CBB cycle attenuation in the light, we pursued two complementary strategies. First, we examined CP12 homologs encoded by cyanophages. The genes for these proteins are expressed in infected hosts during daylight and have been proposed to contribute to host metabolic reprogramming toward a more reduced state during infection ^61,62^. Such phage-derived CP12 variants may exhibit altered redox responsiveness and therefore represent candidates for redox-independent CBB cycle inhibition. Second, we applied rational protein design to engineer synthetic CP12 variants in which the structural stabilization normally mediated by redox-sensitive cysteine bridges was replaced by redox-insensitive interactions. This strategy aimed at generating constitutively structured CP12 variants capable of inhibiting GAPDH and PRK independently of the cellular redox state.

To identify potentially **r**edox-**i**ndependent **CP12** variants (**riCP12**), we screened cyanophage genomes for CP12 homologs using annotation- and motif-based searches. Eighty unique full-length CP12 homologs were identified. All retained the conserved C-terminal GAPDH-binding motif and cysteine pair, whereas the N-terminal cysteine pair and most of the PRK-binding core sequence characteristic of the canonical cyanobacterial CP12-C/N type were absent **(Fig. 4 A,B)**. These phage-derived CP12 proteins correspond to the CP12-C(M) type which is predicted to be less disordered ^63^. AlphaFold-based structural modelling of representative phage CP12 variants further suggested the presence of conserved alternative intramolecular interactions and additional proline residues in the loops that may stabilize the folded conformation independently of redox-controlled disulfide bond formation **(Fig. S2)**. Notably, the residues involved in these predicted interactions were highly conserved among the 80 homologs and differed from the corresponding positions in CP12^6803^ **(Fig. 4 A,B)**. Based on sequence conservation and diversity, we generated an artificial consensus variant (CP12^phage^^1^) and additionally selected two representative CP12 proteins from *Prochlorococcus* phage P-HM2 (CP12^phage^^2^) and *Synechococcus* phage ACG-2014c (CP12^phage^^3^) for functional evaluation **(Fig. 4 B)**.

**Figure 4:**
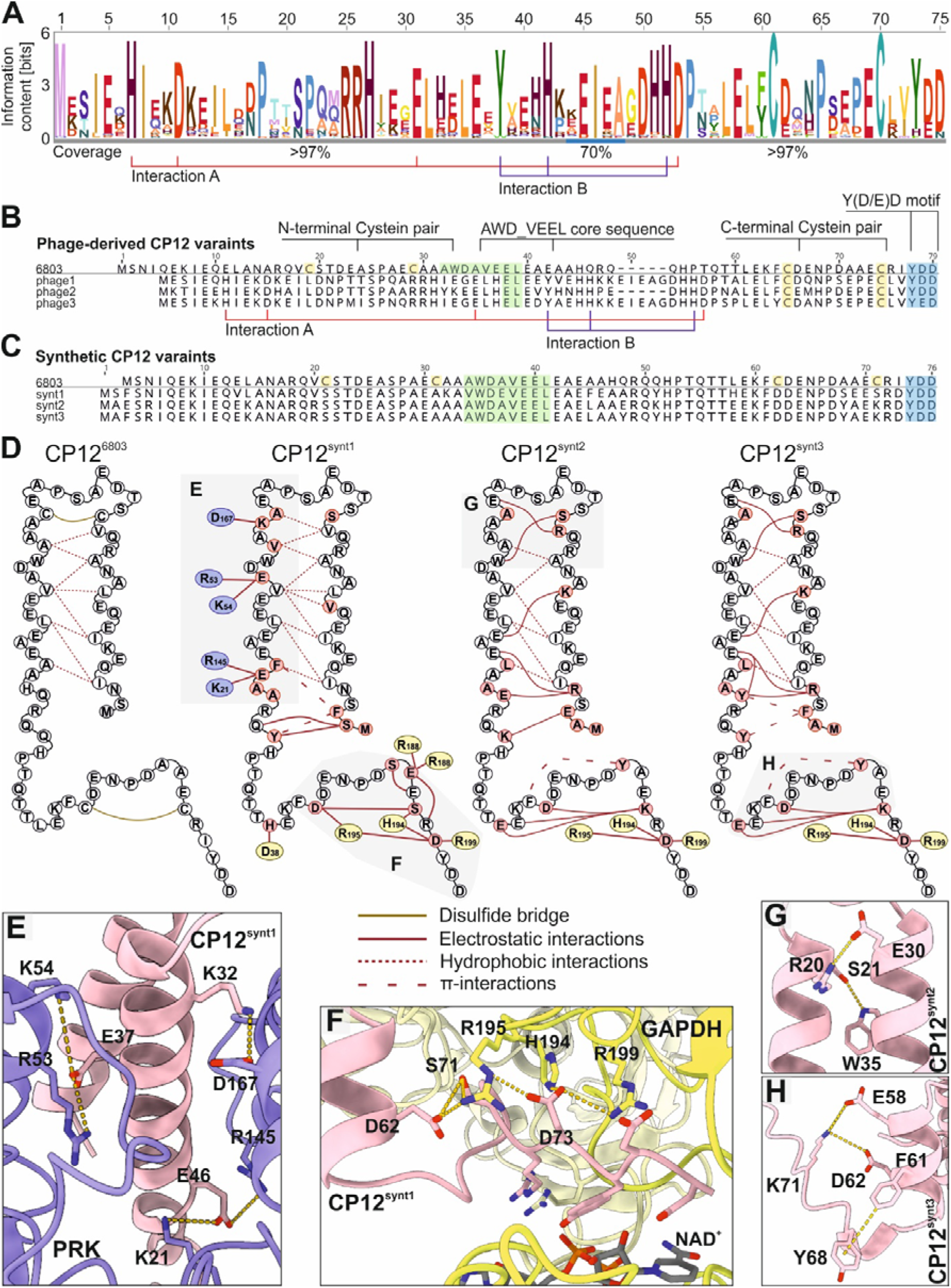
Analysis of phage-derived and design of synthetic potentially redox-insensitive CP12 variants. **A**: HMM logo of 80 unique CP12 proteins from cyanophages. **B:** Alignment of the phage-derived CP12 sequences included in further analysis with the CP12 from *Synechocystis* (6803). Consensus sequence of phage-derived CP12 proteins (phage1); CP12 from *Prochlorococcus* phage P-HM2 (CP12^phage2^); *Synechococcus* phage ACG-2014c (CP12^phage3^). Interactions networks in phage-derived CP12 in AlphaFold predictions absent from CP12 of *Synechocystis* shown in **(Fig. S2)** are indicated in (A) and (B). **C:** Alignment of the designed synthetic CP12 sequences with the CP12 from *Synechocystis* (CP12^6803^). **D:** Schematic representation of the structure of CP12^6803^ and the designed synthetic CP12 variants. Hydrophobic interactions and disulfide bonds stabilizing the structure are shown for CP12^6803^. Mutated positions (red) and their potential intramolecular or intermolecular interactions with PRK (blue) or GAPDH (yellow) are presented for the synthetic variants. Structures of the underlaid regions are displayed in the indicated panels. **E-F:** Homology model of the CP12-GAPDH-PRK complex from *Synechocystis* with mutations to CP12^synt1^. Sidechains of the mutated amino acids and their designated interacting positions are displayed. G-H: AlphaFold models of CP12^synt2^ (G) and CP12^synt3^ (H) highlighting the interactions replacing the N-terminal or C-terminal disulfide bridge of CP12^6803^.

In parallel, based on CP12^6803^, we designed synthetic riCP12 variants. Because canonical CP12 activity depends on redox-controlled disulfide bond formation, simple substitution of the cysteine residues destabilizes and thereby removes redox reactivity of the protein permanently ^36^. We therefore combined replacement of all four cysteine residues with compensatory mutations designed to preserve the structured state required for inhibition of GAPDH and PRK **(Fig. 4C)**. Structure-guided design was performed based on a homology model of the GAPDH–PRK–CP12 complex **(Fig. 4E–H)**. Two complementary strategies were pursued: (i) stabilization of the N- and C-terminal loops that are normally maintained by the redox-sensitive cysteine bridges through introduction of alternative interaction networks, and (ii) reinforcement of protein–protein interfaces with GAPDH and PRK to maintain complex formation independent of the redox state. Three synthetic variants were constructed. The CP12^synt^^1^ variant focused on strengthening intermolecular interactions with GAPDH and PRK to compensate for the loss of cysteine-mediated stabilization. The CP12^synth^^2^ and CP12^synth^^3^ variants instead emphasized intramolecular stabilization of the structured conformation through alternative ionic and aromatic interaction networks **(Fig. 4D)**. All synthetic riCP12 variants, together with the phage-derived riCP12 proteins and CP12^6803^ as a control, were included in subsequent functional analyses.

### Engineered riCP12 variants inhibit GAPDH and PRK in the light thereby inducing a drastic growth arrest and shifting the redox state toward H_2_ production

*In vivo*, a redox-independent, active CP12 variant is expected to attenuate CBB cycle activity immediately upon induction of corresponding gene expression. To test this, we introduced sequences encoding the six selected riCP12 candidates as well as CP12^6803^ on replicative plasmids into *Synechocystis* WT. The *ricp12* genes were put under control of the inducible promoter of the *nrsB* gene that responds to micromolar concentrations of Ni^2+^ ions ^64^. The genetic configurations of all generated mutants as well as their experimental verification is shown in the Appendix **(Fig. S3)**. Autotrophic growth of recombinant *Synechocystis* strains was monitored following induction of gene expression as a functional readout for CBB inhibition. Under standard conditions of low light (50 µmol photons m^-2^ s^-1^) and ambient CO_2_ (0.04% [v/v]), which support only moderate growth rates, induction of *ricp12* gene expression had no detectable effect on growth for any strain (exemplified in **Fig. S4 & S5)**. In contrast, under high light (>150 µmol photons m^-2^ s^-1^) and high CO_2_ supply (4 %[v/v]), several strains carrying the inducible *ricp12* variants – but not *cp12*^6803^ – exhibited in a pronounced growth defect compared to an empty vector strain **(Fig. 5A)**. Upon induction of *ricp12* expression, growth ceased approximately 12 h after inducer addition, and in several cases culture density declined substantially. It should be noted that the slightly reduced growth rates observed in the control strains were attributable to the inducer itself, which showed mild toxicity at the concentrations used consistent with previous reports ^65^. In all cases, however, the growth reduction in strains harboring engineered riCP12 variants was far more pronounced than in the controls. The extent of the growth inhibition varied among the phage-derived riCP12 variants, with the consensus sequence variant (CP12^phage^^1^) exerted the strongest effect. All synthetic riCP12 variants caused an even more pronounced phenotype. Within 24 h after induction, cultures expressing genes for CP12^synth^^1–3^ exhibited bleaching (**Fig. S5G**) accompanied by a decrease in optical density (**Fig. 5A**). Among these variants, CP12^synt^^1^, which was primarily designed to strengthen the protein-protein interfaces with GAPDH and PRK, caused the most severe growth impairment. Remarkably, the magnitude of growth inhibition was titratable with inducer concentration **(Fig. S4, S5)**. It should also be noted that growth arrest was transient: cultures resumed growth 12-24 h after induction **(Fig. 5A)**, which could be partially due to accompanied export of Ni^2+^ ions via the Nrs system ^66^.

**Figure 5:**
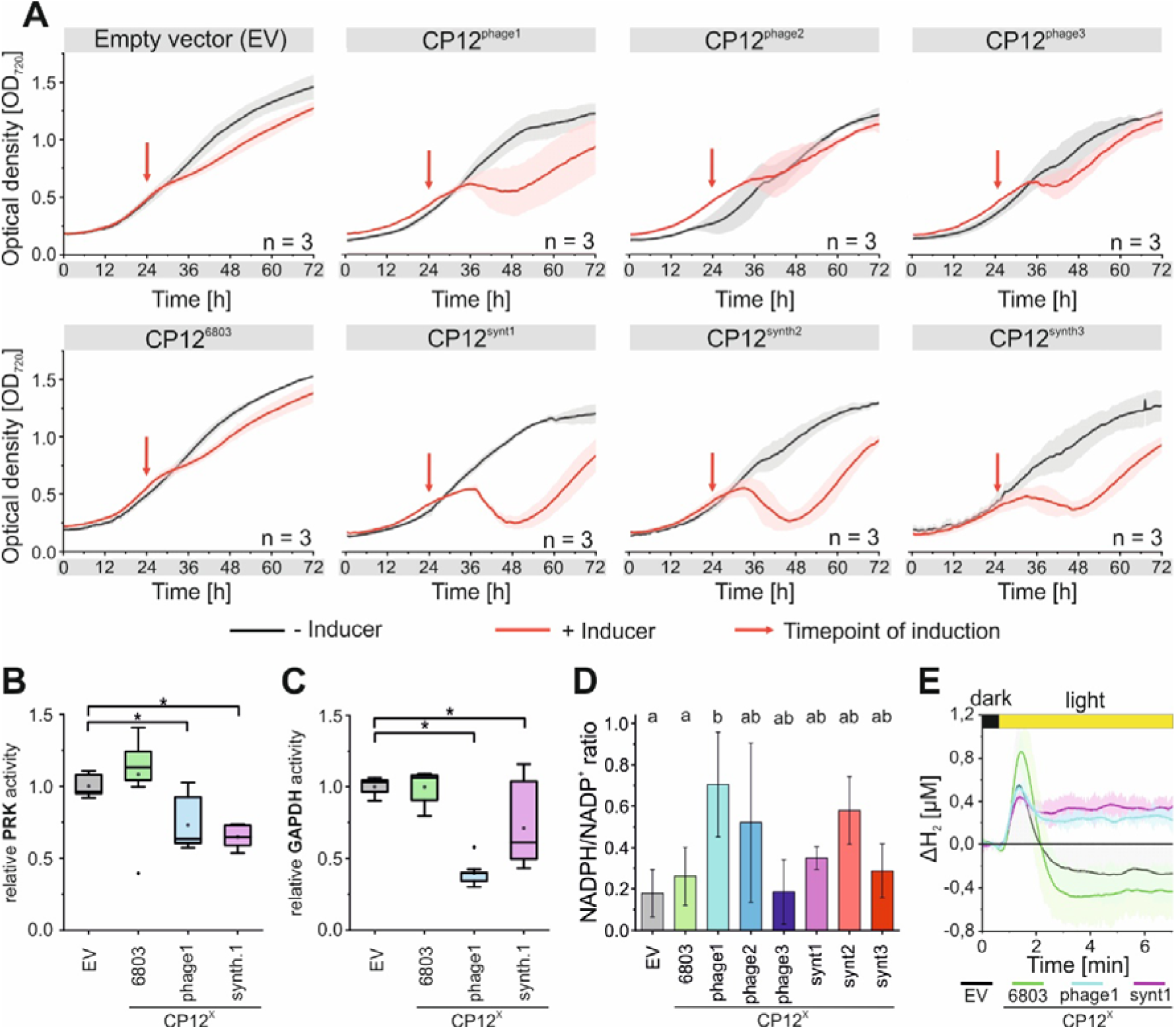
Screening phage-derived and synthetic CP12 variants for redox-independent CBB cycle inhibiting activity. **A**: Growth curves of *Synechocystis* strains expressing *cp12* variants from an inducible promoter with and without induction. After an adaption phase, the strains were grown under 200 µE m-^2^ s-^1^ with 4% CO2. 10 µM NiSO_4_ was added to induce *cp12* expression at t = 24 h (red arrows). Data show the smoothed average from automatic growth monitoring using the MC-1000 ± SD of three biological replicates **B-C:** Enzyme activity of PRK (B) and GAPDH (C) in the crude extract 12 h after induction of *cp12* expression in reduced conditions. Data were normalized to the average activity of the empty vector control strain. The box plots from three biological replicates with three technical replicates each show median (line), average (square), 25 – 75% quartiles (boxes), and minimum and maximum values within 1.5× the interquartile range (whiskers). Statistical significance (p□<□0.05 (*)) between groups was assessed using a Tukey post hoc test. **D:** Reduction state of the NADP pool 24 h after induction of *cp12* expression. Displayed is the average ± SD of three biological replicates and the grouping in a Tukey post hoc test. **E**: H_2_ evolution during dark-light transition in the *Synechocystis* WT expressing ri*cp12* gene variants. Data show the smoothed average of 3 biological replicates ± SD and was baseline adjusted to the average concentration in the last minute before illumination. The measurements were conducted in the liquid phase using UniSense electrodes (full graphs are shown in **Fig. S7**). To maintain anaerobic conditions also under photosynthetic conditions (light phase) cultures were supplemented with 10 mM glucose an enzyme mixture of 40 U glucose oxidase and 50 U catalase (God/Cat). EV, empty vector control.

To directly examine the anticipated function of designed riCP12 variants, we analyzed GAPDH and PRK activities in crude extracts of two representative strains harboring CP12^phage^^1^ and CP12^synth^^1^ in comparison to control strains either overexpressing *cp12*^6803^or harboring an empty vector. Both the synthetic and phage-derived variants significantly reduced the overall GAPDH and PRK activities compared to the controls **(Fig. 5C,D)**. Notably, inhibition was especially observed under reducing conditions, demonstrating redox-independent activity of the *riCP12* variants (**Fig. S6**). GAPDH activity in crude extracts (cell free) from WT and the empty vector control gradually decreased under increasingly oxidizing conditions. In contrast, PRK activity reached minimal levels already under mildly oxidizing conditions. To assess the contribution of CP12, both enzyme activities were also analyzed in a Δ*cp12* strain lacking the chromosomal *cp12* gene. In this background, PRK deactivation under oxidizing conditions was retained, whereas its gradual redox-dependent activation was lost, indicating that the residual inhibition is mediated by intrinsic cysteine residues of PRK^32^. GAPDH activity was generally lower and more variable in Δ*cp12* crude extracts and largely lost its redox responsiveness, except under strongly oxidizing conditions, where activity was abolished in both WT and Δ*cp12* crude extracts (**Fig. S6A,B**). Because the chromosomal *cp12* gene was retained in all *riCP12*-expressing strains, the characteristic redox-dependent regulation of both PRK and GAPDH observed in the WT was also preserved in these strains (**Fig. S6C,D**).

Moreover, to assess the impact of riCP12 variants on cellular redox state, we quantified the NADPH/NADP^+^ ratio 24 h after induction (**Fig. 5B**). Expression of riCP12 variants led to a more reduced NADP pool compared to control strains, consistent with impaired CO_2_ fixation. Interestingly, the magnitude of NADP pool reduction did not strictly mirror the growth phenotype. CP12^phage^^1^ caused the strongest NADP reduction, whereas among the synthetic variants CP12^synt^^2^ induced the largest redox shift. In line with its minor growth effect, CP12^phage^^3^ did not alter the NADP pool relative to controls. Based on the strength and duration of the growth phenotype, strains producing CP12^phage^^1^ (phage-derived) and CP12^synt^^1^ (synthetic) were selected for further characterization as being representative for a redox-insensitive activity.

To assess the impact of riCP12 on H_2_ production, the H_2_ evolution kinetics during a dark-light transition were analyzed in both strains in comparison to respective control strains (**Fig. 5E**). First, the cells were incubated in the dark, where they accumulated H_2_ fermentatively at levels comparable to the WT under anaerobic conditions (**Fig. S7,** compare with **Fig. 1B**). Upon illumination, H_2_ accumulation increased transiently for approximately one minute before declining to levels below those observed before the dark-light transition, consistent with activation of the CBB cycle and restoration of a more oxidized redox state **(Fig. S7)**. Overexpression of *cp12*^6803^ resulted in H_2_ accumulation kinetics similar to an empty vector control strain but with a moderately higher H_2_ peak and enhanced subsequent re-uptake, suggesting subtle alterations in CBB cycle activation due to increased CP12 abundance **(Fig. 5E)**. In contrast, expression of *ricp12* variants markedly altered the H_2_ kinetics after illumination. While the initial H_2_ peak was similar (CP12^phage^^1^) or slightly reduced (CP12^synt^^1^) relative to the control strains, the subsequent decline in H_2_ concentration was significantly attenuated **(Fig. 5E)**. For both *ricp12*-expressing strains, H_2_ levels in the light remained above the pre-illumination value **(Fig. 5E)**. This impaired H_2_ re-uptake supports a shift in the cellular redox balance in the light, consistent with sustained restriction of CBB cycle activity. Collectively, these results demonstrate that both phage-derived and rationally engineered CP12 variants function as redox-independent inhibitors of GAPDH and PRK *in vivo*, enabling inducible attenuation of CBB cycle activity under photosynthetic conditions leading to shifts in the cellular redox-status and thereby supporting H_2_ production in a WT background.

### H_2_ production can be enhanced further by integration of NDH-1 deletion, hydrogenase overproduction, and inhibition of CBB cycle via riCP12

The individual interventions and initial combinations described above uncovered distinct limitations to photosynthetic H_2_ production, including intracellular redox status, catalytic capacity, ATP/NADPH balancing, and competition for electrons. However, these limitations and thereby the effects of the presented interventions were dependent on the applied conditions. To assess combinatorial effects and their dependence on the metabolic conditions, we combined the individual interventions and examined the resulting H_2_ production rates under various conditions. For that purpose, we first introduced the two riCP12 variants CP12^synth^^1^ and CP12^phage^^1^ as well as CP12^6803^ and an empty vector as controls into the previously established M55 mutant and screened the H_2_ production rates under variable irradiation strength, *ricp12* induction condition, and uncoupler concentrations in mixotrophic conditions using the established glucose-dependent oxygen-removal strategy (**Supplementary Table 1**). Thereby, the uncoupler DNP had a similarly positive effect on H_2_ production for the riCP12 and CP12^6803^ carrying strain as demonstrated for the initial M55 strain carrying an empty vector (**Fig. 1D, S8**). Regardless, the maximal H_2_ production rate of 0.05 µmol min^-1^ L^-1^ OD_750_^-1^, which was achieved under high light conditions, was not increased by the introduction of the riCP12’s. However, the strains expressing a *ricp12* genes exhibited H_2_ production rates close (up to 80 %) to the observed maximal rates under high light conditions already under conditions of low light intensity (25 µE m^-2^ s^-1^) limiting photosynthetic electron supply. In comparison, the empty vector control strain evolved H_2_ only at rates much lower than the maximal rate (∼ 10 %) (**Fig. S8**). Combined with the achieved drastic increase of H_2_ formation by hydrogenase overproduction in the NDH-1 deletion strain (**Fig. 2**), this emphasizes the hydrogenase capacity as the main limiting factor in this strain. The observed differences under restricted photosynthetic activity, however, suggest that redistribution of electrons can be achieved with riCP12.

To assess the effect of electron redistribution by riCP12 under conditions of higher hydrogenase capacity, we finally incorporated CP12^synt^^1^, the riCP12 variant that caused the strongest effects before, into the Δ*ndhB* Δ*hox* P_J23119_*:hoxEFUYH* strain. The resulting strain Δ*ndhB* Δ*hox* P_J23119_*:hoxEFUYH* P*_nrsB_*_*cp12*^synth^^1^ as well as its predecessor without riCP12, again, were analysed for their H_2_ formation rates under autotrophic conditions using the co-culture assay with and without pmf-uncoupling, different regimes for the induction of *cp12*^synth^^1^, and varying light intensities.

The inducer NiSO_4_, which simultaneously serves as co-factor for the hydrogenase, did not considerably alter the H_2_ production rate of the Δ*ndhB* Δ*hox* P_J23119_*:hoxEFUYH* strain **(Fig. 6A)**. Moreover, in contrast to the mixotrophic conditions neither reduction of the light intensity in the production phase to 25 µE m^-2^ s^-1^ nor its increase to 200 µE m^-2^ s^-1^ had a significant effect under these conditions **(Fig. 6B)**. Conversely to the strongly enhancing effect of pmf-uncoupling by DNP under mixotrophic conditions and the considerable increase under the same conditions employed here to the H_2_ production of the M55 mutant **(Fig 1D)**, addition of DNP to both the hydrogenase overproducing strains with and without riCP12 caused a reduction of their productivity across conditions by 25 % and 27 %, respectively **(Fig. 6C,G)**. In direct comparison with Δ*ndhB* Δ*hox P_J23119_::hoxEFUYH*, the strain additionally encoding CP12^synth^^1^ showed a 94 % increased mean H_2_ formation rate **(Fig. 6D)**. This suggests a supportive effect of CBB cycle attenuation on continuous H_2_ formation given sufficient hydrogenase capacity. Moreover, H_2_ production rates of Δ*ndhB* Δ*hox* P_J23119_*:hoxEFUYH* P*_nrsB_*_*cp12*^synth^^1^ were further increased by 38 % upon raising the light intensity from 10 to 50 µE m^-2^ s^-1^ **(Fig. 6F)**. In contrast, inducible modulation of the CP12^synth^^1^ effect was modest and condition-dependent. While consistent trends were observed across experimental conditions, these effects did not reach statistical significance in the global model. Notably, maximal induction of *ricp12* even reduced H_2_ production **(Fig 6E)**. Under otherwise favourable conditions, a moderate induction of *cp12*^synth^^1^ expression noticeably increased observed rates with a time-resolved trend consistent with CP12^synth1^’s effect on growth in the *Synechocystis* WT and resulted in a maximal observed H_2_ production rate of 0.96 µmol min^-1^ L^-1^ OD_750_^-1^.

**Figure 6:**
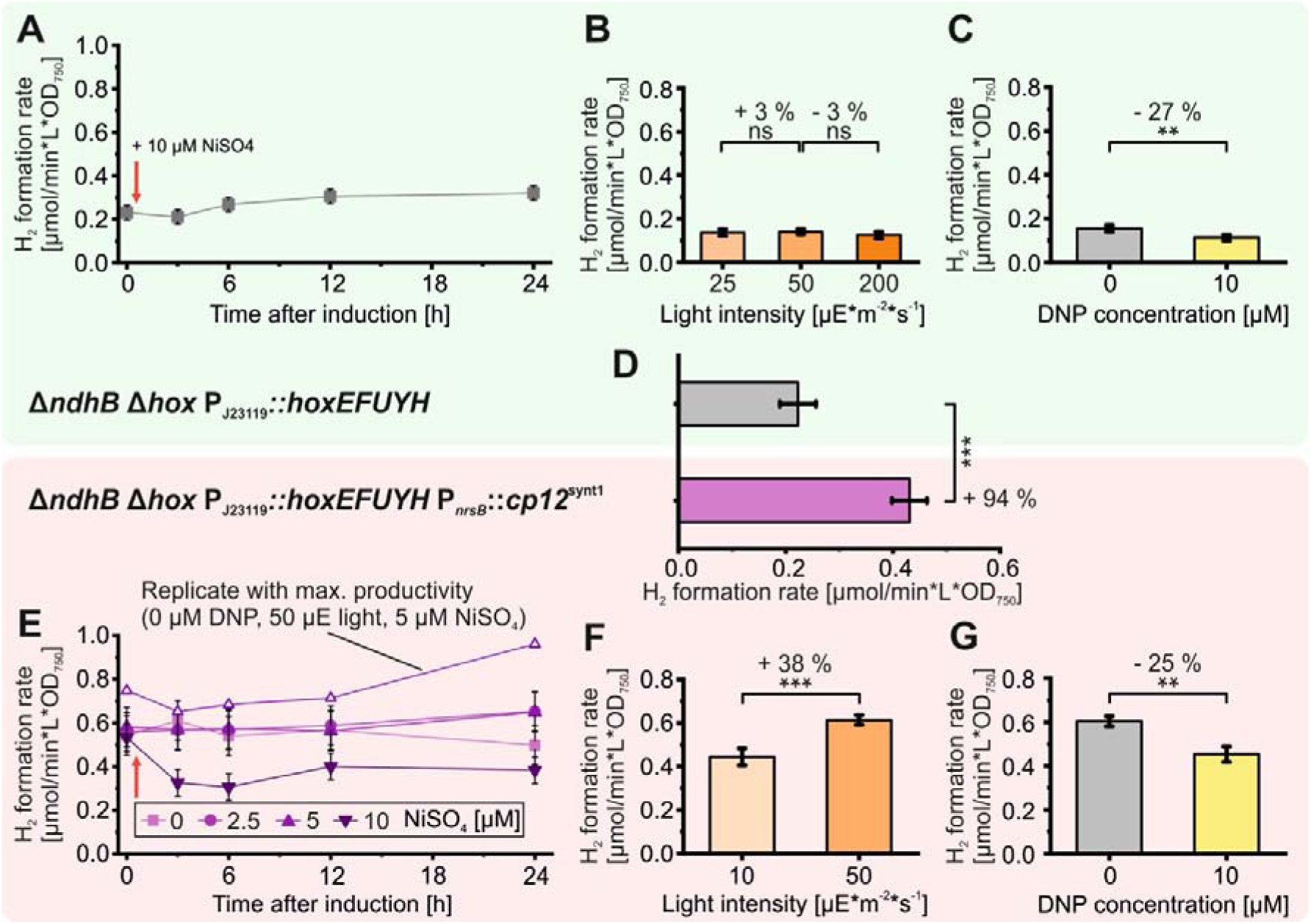
Integration of factors influencing H_2_ production of hydrogenase overproducing, NDH-1-deficient *Synechocystis strains*. The *Synechocystis* Δ*ndhB* Δ*hox PJ23119::hoxEFUYH* strain (A-D) or a strain additionally expressing *cp12*^synth1^ from a Ni^2+^-inducible promoter (D-G) were testes for their H_2_ production rates in *Pseudomonas* co-cultures fed with 10 mM citrate under various conditions. Production rates were calculated from H_2_ concentrations in headspace of sealed cultures quantified by GC after ∼ 24 h. Data were analyzed using linear mixed effect models (LMM) and estimated marginal means (EMM) are displayed. A, E: Effect of addition (red arrows) of the inducer for *cp12*^synth1^-expression (NiSO_4_) to cultures without (A) or with (E) *cp12*^synth1^ encoded. Data with open triangles represent the single replicate with the highest H_2_ production rate while closed symbols display EMM. B, F: Effect of light intensity during the H_2_ production assay with the strains without (B) or with (F) *cp12*^synth1^ encoded. C, G: Effect of DNP addition to the H_2_ production assay with the strains without (C) or with (G) *cp12*^synth1^ encoded. D: Comparison of the strain without to the strain with *cp12*^synth1^. Statistical significance (p□<□0.05 (*); <□0.01 (**); <□0.001 (***)) between groups was assessed using post hoc pairwise comparisons based on the LMM with appropriate correction for multiple testing.

Although variability was initially reduced by overexpressing the *hoxEFUYH* operon from a constitutive promoter **(Fig. 2G)**, the Δ*ndhB* Δ*hox* P_J23119_*::hoxEFUYH* strain and especially its *cp12* expressing derivates still exhibited considerable variation between independent experiments **(Fig. 2G**, **Fig. 6)**. Moreover, variability between clones and the number of clones losing their H_2_ production capacity over time after initial verification of their genetic background and H_2_ production rate, increased with the number of interventions, suggesting emerging instability under these highly perturbed conditions. Resequencing of the respective plasmids revealed mutations especially connected to overexpression of the *hox* operon **(Fig. S9)**.

## Discussion

Previous studies identified several constraints on cyanobacterial H_2_ formation and proposed engineering strategies to alleviate them individually ^11,38,40,45,47^. Here, we integrated these approaches with newly developed interventions optimized for sustained photoautotrophic H_2_ production, allowing us to systematically dissect the constraint architecture governing long-term H_2_ evolution. Rather than identifying a single universal bottleneck, successive interventions shifted the physiological state of the cell and exposed new limitations, demonstrating that sustained H_2_ production is governed by a hierarchical and state-dependent network of metabolic constraints.

The first prerequisite for sustained H_2_ evolution is thermodynamic feasibility. Because proton reduction by the bidirectional hydrogenase operates close to thermodynamic equilibrium under physiological conditions, sustained flux requires highly reduced ferredoxin and NADPH pools. Cellular redox poise is therefore determined by the balance between photosynthetic electron transport and competing electron sinks, including the CBB cycle, cyclic electron transport, and oxygen reduction. Accordingly, anaerobiosis not only preserves hydrogenase activity but also suppresses oxygen-dependent electron dissipation (e.g., Mehler-like reactions), promoting a more reduced intracellular redox state^45^. Consistent with this model, partial deletion of ferredoxin–NADP^+^ reductase (FNR) affected H_2_ formation dynamics^47^, yet complete deletion was not feasible due to the enzyme’s central metabolic role. Moreover, as reported here the attenuation of CO_2_ fixation through redox-independent CP12 variants (riCP12) also increased the H_2_ reuptake threshold following dark-light transitions similar to partial FNR mutants, indicating a more reduced steady-state redox environment. However, neither intervention alone supported sustained H_2_ formation. In contrast, NDH-1-deficient strains maintained highly reduced redox pools and continuous H_2_ evolution despite an intact CBB cycle, identifying cyclic electron transport as a major competing sink, although additional effects on the CO_2_-concentrating mechanism cannot be excluded^38^.

Once thermodynamic constraints were relieved, ATP/NADPH balancing emerged as the next major limitation under mixotrophic conditions as chemical uncoupling revealed a direct connection between the proton motive force (pmf) and H_2_ production in an NDH-1-deficient mutant **(Fig. 1)**. Photosynthetic linear electron transport generates reducing equivalents and the proton motive force (pmf) in fixed stoichiometry, whereas cyclic electron transport adjusts the ATP/NADPH ratio by increasing ATP generation without net reductant formation. Because cyanobacteria lack an intrinsic mechanism to dissipate pmf independently of electron flow, excessive pmf feeds back to restrict photosynthetic electron transport^53^. Chemical uncoupling demonstrated that this bioenergetic constraint is strongly dependent on the metabolic state of the cell. Under mixotrophic conditions, substrate-level phosphorylation likely reduced the demand for photosynthetically generated ATP, allowing pmf to accumulate and limiting electron throughput ^72^. Dissipation of pmf therefore markedly stimulated H_2_ formation. In contrast, under fully autotrophic anaerobic conditions ATP generation depended almost exclusively on linear electron transport, where ATP demand typically exceeds ATP supply^73^. Under these conditions, pmf buildup and thereby electron throughput was no longer limiting and hydrogenase capacity instead became the dominant constraint.

Consistent with this interpretation, ectopic overexpression of the *hox* operon produced the largest increase in H_2_ evolution, whereas increasing light intensity or attenuating the CBB cycle by riCP12 in the NDH-1 deletion strain alone had little effect. Conversely, the weak stimulatory effect of DNP under autotrophic conditions in the parental M55 strain -but not in hydrogenase-overproducing strains-suggests that DNP may additionally enhance hydrogenase expression under native regulation. Once hydrogenase capacity was increased, however, electron allocation became limiting. Under these conditions, attenuation of the CBB cycle by riCP12 substantially enhanced H_2_ production, demonstrating that competition for photosynthetically derived electrons can become a major constraint even in NDH-1 deletion strains with highly reduced redox pools. Strong induction of riCP12, however, decreased rather than increased H_2_ evolution, likely reflecting excessive suppression of carbon metabolism impairing cellular maintenance, although toxicity of the NiSO_4_ inducer may also contribute^66^. These observations further suggest that ATP/NADPH balancing may again become limiting also under autotrophic conditions when progressively larger fractions of photosynthetic electrons are redirected toward H_2_ production, highlighting inducible ATP-dissipating strategies^74^ as an important target for future metabolic engineering.

Finally, our results identify evolutionary robustness as an additional constraint that became apparent while metabolic limitations have been relieved. Sustained diversion of photosynthetic electrons toward H_2_ was consistently counteracted by metabolic and genetic adaptation. Induction of riCP12 variants caused transient chlorosis and growth arrest followed by recovery within approximately 24 h, indicating rapid network-level compensation, e.g. downregulation of ri*cp12* gene expression or upregulation of *gap2* or *prk* coding for the targeted enzymes. More strikingly, genetic instability became especially apparent when combining hydrogenase overproduction with the expression of ri*cp12* gene variants with strong selection against elevated hydrogenase activity. During construction of strains carrying a single plasmid encoding both *hoxEFUYH* and ri*cp12* for subsequent overexpression, the majority of tested clones obtained after transformation did not carry the intended plasmid. Moreover, clones initially confirmed to carry the intended plasmid and exhibiting high H_2_ production rates often lost this capacity after a few repeated cultivations. Conversely, in strains in which *hoxEFUYH* overexpression and ri*cp12* expression was separated onto two different replicative plasmids and introduced sequentially, most clones carried both plasmids but revealed mutations interrupting *hoxF* and abolishing H_2_ production **(Fig. S9).** Importantly, cultures were maintained under aerobic conditions during strain construction, excluding H_2_ evolution as the direct cause of selective pressure. Notably, repeated emergence of independent mutations in *hoxF* (encoding the diaphorase subunit) suggests strong selective pressure against hydrogenase overproduction itself, which could be counteracted by more balanced and controlled expression of the individual subunits. Moreover, it supports emerging evidence for additional physiological roles of the diaphorase module ^75^.

Collectively, our results define a hierarchical, dynamic, and physiological state-dependent constraint architecture governing sustained photosynthetic H_2_ formation. Thermodynamic feasibility establishes the prerequisite for H_2_ evolution, whereas ATP/NADPH balancing, hydrogenase capacity, and competition for electrons become limiting according to the metabolic state of the cell. In addition, evolutionary robustness must be considered to achieve stable H_2_ formation rates. Importantly, these constraints primarily affect different properties of the system: hydrogenase capacity determines maximal production rates, whereas ATP/NADPH balancing, redox homeostasis, and electron allocation govern the establishment and maintenance of a stable productive state. By using H_2_ evolution as a quantitative systems-level reporter of photosynthetic energy metabolism, our study reveals fundamental principles of redox regulation and energy homeostasis that extend beyond H_2_ formation itself. More broadly, our findings explain why optimization of individual components has yielded only incremental improvements and suggest that future advances will require coordinated engineering of interconnected metabolic and regulatory processes while balancing productivity against long-term cellular fitness.

Beyond defining the hierarchical constraints governing sustained H_2_ production, this study establishes engineered small regulatory proteins as a versatile tool for reprogramming cellular metabolic states and redirecting metabolic fluxes. The effects of riCP12 variants on both growth and H_2_ formation demonstrate that they dynamically remodel carbon metabolism and redistribute photosynthetic electrons toward target reactions, analogous to the proposed function of auxiliary metabolic proteins during cyanophage infection^61,62^. By applying this regulatory engineering strategy, we achieved metabolic reprogramming comparable to partial FNR deletion while avoiding permanent disruption of an essential enzyme and its associated constitutive fitness costs. More broadly, our work highlights regulatory proteins as an underexplored engineering layer that complements conventional enzyme- and pathway-based metabolic engineering. Although phage-derived proteins have long served as indispensable tools in biotechnology, particularly for gene expression^67^, their auxiliary metabolic proteins have rarely been exploited as programmable regulators of metabolism in biotechnological production strains. Our results demonstrate that rational engineering of regulatory proteins -whether inspired by phage proteins or endogenous host factors-can reshape metabolic states without directly altering catalytic capacity. Together with recent advances in protein structure prediction and *de novo* protein design^68–71^, this strategy opens new opportunities to create synthetic regulatory interactions for enzymes that lack natural regulatory partners, substantially expanding the toolbox for programmable metabolic control.

Overall, our findings indicate that future improvements in photosynthetic H_2_ formation are unlikely to arise from optimization of individual enzymes or pathways, but instead require coordinated manipulation of multiple hierarchical constraints while maintaining cellular fitness. More broadly, establishing H_2_ evolution as a systems-level reporter of photosynthetic energy metabolism provides a conceptual framework for dissecting complex metabolic networks and identifying new regulatory strategies to balance metabolic performance with long-term stability.

## Material and Methods

### Strains and growth conditions

The WT of *Synechocystis* was obtained from the Pasteur Culture Collection of Cyanobacteria (PCC). By default, *Synechocystis* strains were grown in shaking flasks with BG11 medium^76^ that was modified as previously described^77^, at 30 °C, 50 µE m^-2^ s^-1^ light intensity, and ambient CO_2_. For strain construction, respective plates (1.5 % BD DIFCO Bacto-Agar) were incubated at a lower light intensity of 25 µE m^-2^ s^-1^. For maintenance of the M55 mutant and other NDH-1 deficient strains, the CO_2_ concentration was increased to 2 % for liquid cultures or 0.2 % for agarose plates, respectively. Media were supplemented with the required antibiotics at the following concentrations (µg ml^-1^; standard cultivation / initial selection): kanamycin (50/10), chloramphenicol (15/5), gentamicin (4/2), spectinomycin (50/10). Initial selections were performed using only the respective antibiotic, while reduced concentrations of all respective antibiotics (25 µg ml^-1^ kanamycin, 10 µg ml^-1^ chloramphenicol, 2.5 µg ml^-1^ gentamicin) were used for *Synechocystis* strains carrying several resistances during standard cultivation. Cultivation of *Synechocystis* in Multicultivator MC-1000 column reactors (Photon System Instruments) was employed for growth analysis with automated density measurements. *E. coli* TOP10 for plasmid construction were cultivated aerobically in Lysogeny Broth (LB) medium in liquid culture or on respective agarose-solidified plates (1.5 % BD DIFCO Bacto-Agar) at 37 °C or, when harboring a pSOMA-based plasmid, at 30 °C. *Pseudomonas taiwanensis* VLB120 was grown aerobically at 30 °C either in LB or M9 medium^78^ that was supplemented with 0.1 % (v/v) US trace element solution^79^ and 2 mM MgSO_4_ as described previously^80^ and 40 mM trisodium citrate instead of glucose as the sole carbon source.

### Plasmid and strain construction

Deletions or interruptions of chromosomal genes in *Synechocystis* were accomplished by homologous recombination and subsequent selection on the respective antibiotic until the strain exhibited full segregation as described before ^81^. Replicative plasmids were inserted into *Synechocystis* via electroporation as described previously ^2,82^. Polymerase chain reaction (PCR) with the GoTaq^®^ PCR master mix (Promega) was employed for verification of successful transformation and segregation. A previously described Golden-Gate cloning (GGC) system ^83^ was used for construction of plasmids. New parts were prepared for integration into the GGC system by PCR with Q5^®^ polymerase (New England Biolabs), or, in case of the *cp12* coding sequences, were ordered as linear double-stranded DNA fragments (Eurofins genomics) with respective overhangs **(Table S3)**. Plasmid construction was carried out in *E. coli* TOP10 following routine protocols. Plasmid sequences were verified via Sanger sequencing (Genewiz). Re-sequencing of plasmids was done via whole-plasmid sequencing (Genewiz). Geneious prime (Dotmatics) was used for plasmid design and analysis of sequencing reads. Details on the employed and generated primers, plasmids, and strains are given in **Supplementary Tables S1-4**.

### H_2_ production assays

For quantification of H_2_ production, samples of *Synechocystis* cultures were taken from the respective cultivation and concentrated by centrifugation at 3,900 xG. Subsequently, pellets were dissolved in BG11 and the OD_750_ was set to 0.5 – 3 in the assay depending on the experiment. Anaerobiosis was ensured by adding glucose or citrate to a final concentration of 10 mM and either an enzyme mix of glucose oxidase and catalase to a final concentration of 40 U mL^-1^ and 50 U mL^-1^ respectively, or *Pseuodomonas* cells concentrated in BG11 medium to a final OD_600_ of 1. Optionally, an aqueous 10 mM NiSO_4_ solution or a solution of 2,4-dinitrophenol in dimethyl sulfoxide (10 mM) was added to the cell suspension to the given concentration. Dissolved H_2_ was quantified using a Clark-type H_2_ electrode (Unisense) in closed glass vials with a total volume of 1.5 mL which was filled completely with the cell suspension. Before the measurement, oxygen was removed from the cell suspension by purging with nitrogen. Initially, cells were kept in darkness for 5 to 10 minutes. Then the vial was illuminated with white light of 200 µE m^-2^ s^-1^. Throughout the measurement, the cell suspension was mixed by magnetic stirring. The robust and sustained H_2_ production by the M55 mutant enabled quantification of H_2_ accumulation in the headspace of sealed cultures by gas chromatography. For that, a volume of 3 mL of the assay suspension was filled into 10 mL glass GC vials which was then crimped with a butyl lid and purged for >5 min with a mix of 90 % nitrogen and 10 % carbon dioxide. Subsequently, the vials were incubated at 30 °C in a shaking incubator with white light (50 µE m^-2^ s^-1^ by default or as given in the figure description). The H_2_ and oxygen concentrations in the headspace were analysed using a Trace 1310 gas chromatograph (ThermoScientific) equipped with a TriPlus RSH Headspace autosampler, a TracePLOT TG-Bond MSieve 5A (30 m x 0.32 mm x 30 µm) column, and a Thermal Conductivity Detector (TCD) with the following settings: 100 µL sample volume, 40 °C syringe temperature, 100 °C inlet temperature, split flow of 20 ml min^-1^ argon carrier gas with a split ratio of 4, constant GC oven temperature of 75 °C, and 100 °C TCD temperature. Linear calibration curves for both gases in the range of 100 ppm to 2 % for H_2_ and 1 % to 20.9 % for oxygen were determined using defined gas mixtures (Air products).

### Enzyme activity assays

Crude extracts were prepared essentially as described previously ^84^. *Synechocystis* cultures (30 ml, OD_750_ = 1–2) were harvested by centrifugation (4,500 × g, 10 min, 4 °C). Cell pellets were resuspended in assay-specific crude extract buffer and disrupted using approximately 200 µL of 0.5 mm glass beads in a Precellys^®^ Evolution Touch homogenizer (Bertin) with five cycles of 30 s agitation followed by 45 s cooling to 4 °C. Cell debris and glass beads were removed by centrifugation (11,000 *g*, 15 min). The supernatant was collected as crude extract and supplemented with 10 % glycerol. Crude extract buffers contained 5 mM L-cysteine, 0.1 mM NAD^+^, and 1 mM phenylmethylsulfonyl fluoride (PMSF) in Milli-Q® water and differed in their buffering system and Ethylenediaminetetraacetic acid (EDTA) concentration. For PRK assays, the buffer consisted of 15 mM tris(hydroxymethyl)aminomethane (Tris) and 0.4 mM EDTA (pH 8.0), whereas for GAPDH assays it consisted of 50 mM K_2_HPO_4_/KH_2_PO_4_ and 1 mM EDTA (pH 7.5). For redox treatments, crude extracts were incubated with equal volumes of either dithiothreitol (DTT) or oxidized glutathione at various concentrations for 20 min at 30 °C.

PRK activity was determined using a coupled enzymatic assay adapted from ^85^. Ribulose-5-phosphate (Ru5P) was generated in situ by incubating 1 mM ribose-5-phosphate with 1 U ribose-phosphate isomerase in 50 mM bicine buffer (pH 8.0) containing 10 mM MgCl_2_, 40 mM KCl, and 0.5 mM EDTA for 20 min at 30 °C, followed by heat inactivation (95 °C, 10 min). PRK activity was measured in a 100 µl reaction containing 1 mM phosphoenolpyruvate, 0.15 mM NADH, 0.5 mM ATP, 1 U pyruvate kinase, 2.75 U lactate dehydrogenase, and 15 µL crude extract. The reaction was initiated by addition of 48 µL Ru5P mixture. Oxidation of NADH was monitored fluorometrically (excitation 340 nm, emission 460 nm) in opaque black flat microtiter 96-well-plates (Nunc) using an Infinite 200 PRO plate reader (TECAN). Negative controls lacking Ru5P were subtracted from sample measurements, and positive controls containing ADP verified that the coupling enzymes were not rate-limiting. Activities were normalized to the protein content of the crude extracts.

GAPDH activity was measured using a fluorescence-based assay adapted from previously described approaches ^86,87^. The substrate 1,3-bisphosphoglycerate (BPGA) was generated by preincubating 33 mM ATP, 66 mM glycerate-3-phosphate, and 4.5 U phosphoglycerate kinase in the presence of MgCl_2_ and EDTA for 20 min at 30 °C. Enzyme activity was measured in a 100 µl reaction containing 50 mM glycylglycine buffer, 10 mM MgCl_2_, 40 mM KCl, 0.5 mM EDTA, 0.25 mM NADPH, 15 µL crude extract, and 15 µL BPGA mix. The decrease in NADPH fluorescence was monitored as in PRK assay. Negative controls lacking BPGA were subtracted from all measurements. Positive controls using rabbit muscle GAPDH and NADH verified substrate availability. Activities were normalized to the protein concentration of the crude extracts.

### Analysis of NAD(P)(H) pool

For the analysis of intracellular NAD(P)(H) pools, *Synechocystis* cultures were grown in BG11 medium in biological triplicates (35 mL) starting from an initial OD_750_ of 0.05. Cultures were grown under standard cultivation conditions and expression of the *cp12* genes was induced after two days by addition of 10 μM NiSO_4_. Cells were harvested one day after induction. Cell densities were determined by measuring OD_750_ prior to sampling. For metabolite extraction, 10 mL of culture per replicate were rapidly (approx. 10 s) transferred to 15 mL centrifuge tubes and snap-frozen in liquid nitrogen immediately. Samples were subsequently lyophilized to remove water. Lyophilized cells were transferred to 2 mL Precellys tubes containing 200 μL 0.5 mm glass beads. Metabolites were extracted by adding 500 μL extraction buffer (acetonitrile:methanol:water, 40:40:20) containing 0.1 M formic acid that was precooled to -20 °C. The extraction buffer was purged with nitrogen prior to use and tubes were flushed with nitrogen before sealing to minimize oxidation of redox-sensitive metabolites. Cells were disrupted stream in five cycles of 30 s at 10,000 *g* with 45 s pauses using a Precellys^®^ Evolution Touch homogenizer (Bertin) that was kept at −40 °C to −50 °C via a liquid-nitrogen cooled air. Cell debris and beads were removed by centrifugation (30 min, 17,000 *g*, −9 °C) and the supernatant was used for analysis. Extracts were either analysed immediately or stored at −80 °C for less than 24 h prior to measurement. Metabolite analysis was performed by ion chromatography–mass spectrometry (IC-MS) using a Dionex™ ICS-6000 HPIC system equipped with a IonPac AS11-HC-4 µm 2×250 mm column coupled to an Orbitrap Exploris 240 mass spectrometer (both ThermoScientific) equipped with an electrospray ion source (H-ESI). IC settings: The gradient of the KOH eluent generated from a EGC 500 KOH cartridge was set to 1 mM for 5 min, increased linearly to 20 mM until 23 min, followed by a steeper gradient to 80 mM at 38 min, held for 2 min, and then returned to 1 mM for re-equilibration; current of an ADRS_2 mm suppressor was initially set to 10 mA, increased to 20 mA after 16 min, 30 mA after 25 min, 47 mA after 30 min, 64 mA after 35 min, and 76 mA after 38 min before being gradually reduced to 5 mA with the decreasing eluent concentration; isocratic makeup flow of 0.15 ml min^-1^ methanol; isocratic regenerant flow of 0.5 ml min^-1^ water; isocratic eluent flow of 0.38 ml min^-1^ water; autosampler cooled to 4 °C; 10 µL sample loop volume with a factor 3 loop overfill; 30°C column temperature; conductivity detector was kept at 25 °C with a 5 Hz data collection rate; analyte was diverted to the MS starting from 21 min after injection. MS settings: 2500 V at spray tip in negative ion mode; 50 arb units sheath gas flow; 10 arb unit auxiliary gas flow; 325 °C ion transfer tube temperature; 300 °C vaporizer temperature. Target compounds were ionized in negative mode and detected using the following mass windows: detection ±1 m/z; quantification ±0.01 m/z: NAD^+^ - 540.0502, NADH – 331.55518, NADP^+^ - 309.50452, and NADPH – 371.5354. Linear calibration curves in the concentration range of 5 to 1,000 nM were generated using metabolite standards prepared in extraction buffer containing BG11 salts at concentrations matching the samples.

### In silico sequence and structure analysis

Sequence analyses and *in silico* cloning were performed using Geneious Prime 2021.2.2 (Biomatters). Multiple sequence alignments were generated with Clustal Omega v1.2.2 ^88^, and pairwise protein alignments were calculated using the Geneious Prime alignment tool with the BLOSUM62 substitution matrix ^89^. Sequence information for CP12, PRK, and GAPDH proteins was obtained from the NCBI GenBank database using genome searches and BLAST ^90^. The profile hidden Markov model (HMM) was prepared using the skylign webservice ^91^.

Homology models were generated using the SWISS-MODEL server ^92^ with default settings. The structure of the *Synechocystis* sp. PCC 6803 CP12–GAPDH–PRK complex was constructed from individual homology models of CP12, GAPDH, and PRK using the corresponding complex from *Thermosynechococcus vestitus* BP-1 (PDB: 6GVE) as template ^32^. The modeled proteins were structurally aligned to their respective template counterparts to assemble the complex, and the position of the NAD^+^ ligand was adopted from the template structure based on the high conservation of the GAPDH sequence. Protein structure visualization and structural analysis were performed using UCSF ChimeraX ^93^. Structural predictions for phage-derived CP12 variants were generated using AlphaFold3 ^68^.

### Statistical analysis

Standard statistical analyses are described in the corresponding figure legends. For experiments comparing strains across multiple environmental conditions, linear mixed-effects models were fitted in R (version 4.5.3) using the **lme4** package, with strain (or genotype) specified as a fixed effect and experimental condition as a random intercept. Experimental conditions were defined as matched comparisons performed under identical assay conditions (including light intensity, DNP treatment, assay duration, and sampling time where applicable), allowing variation between conditions to be accounted for while estimating the overall strain effect. For induction experiments, data were analysed using a linear mixed-effects model with induction level, sampling time, assay conditions, and their interactions included as fixed effects. Estimated marginal means (EMMs) were calculated using the emmeans package and are shown throughout the figures. Pairwise comparisons between groups were performed using post hoc contrasts of the fitted models with Tukey adjustment for multiple testing. Degrees of freedom for mixed-effects models were estimated using the Satterthwaite approximation implemented in the lmerTest package. Statistical significance was defined as *P* < 0.05.

## Acknowledgements

We acknowledge the use of the facilities of the Centre for Biocatalysis (MiKat) and H_2_Saxony at the Helmholtz Centre for Environmental Research – UFZ. We thank the current and former members of the involved departments as well as Marius Theune, Jens Appel, and Kirstin Gutekunst (University of Kassel) for fruitful discussions and experimental support. Furthermore, we appreciate the excellent technical assistance provided by Daniel Spindler. We declare the use of AI tools (ChatGPT 5.5) to polish grammar and clarity and support coding for statistical analysis.

## Author contributions

M.A.I. and S.K. conceived the study, designed experiments, interpreted the data, and wrote the manuscript. M.A.I. designed riCP12 proteins, constructed most strains, and performed most experiments. A.S. and S.G. designed the co-culture assay to analyse H_2_ formation. S.G. and J.W. performed experiments comparing the co-culture assay to the established enzymatic oxygen-removal assay. Supervised by M.A.I., L.K. performed the GAPDH and PRK activity assays. F.B. contributed to the design of the study, strain construction and verification. R.S. supported strain construction and H_2_ analytics. P.W.W., M.A.I., and J.O.K. established and performed the analysis of the redox-carrier pools. A.S. and S.K. supervised and managed the project. A.S., J.O.K., and S.K. acquired funding. All authors contributed to data analysis, reviewed and edited the manuscript, and approved the final version.

