## Appendix for "Multilevel engineering of cyanobacterial energy metabolism advances photosynthetic hydrogen production while revealing its constraints"

### Supplementary Figures and Tables

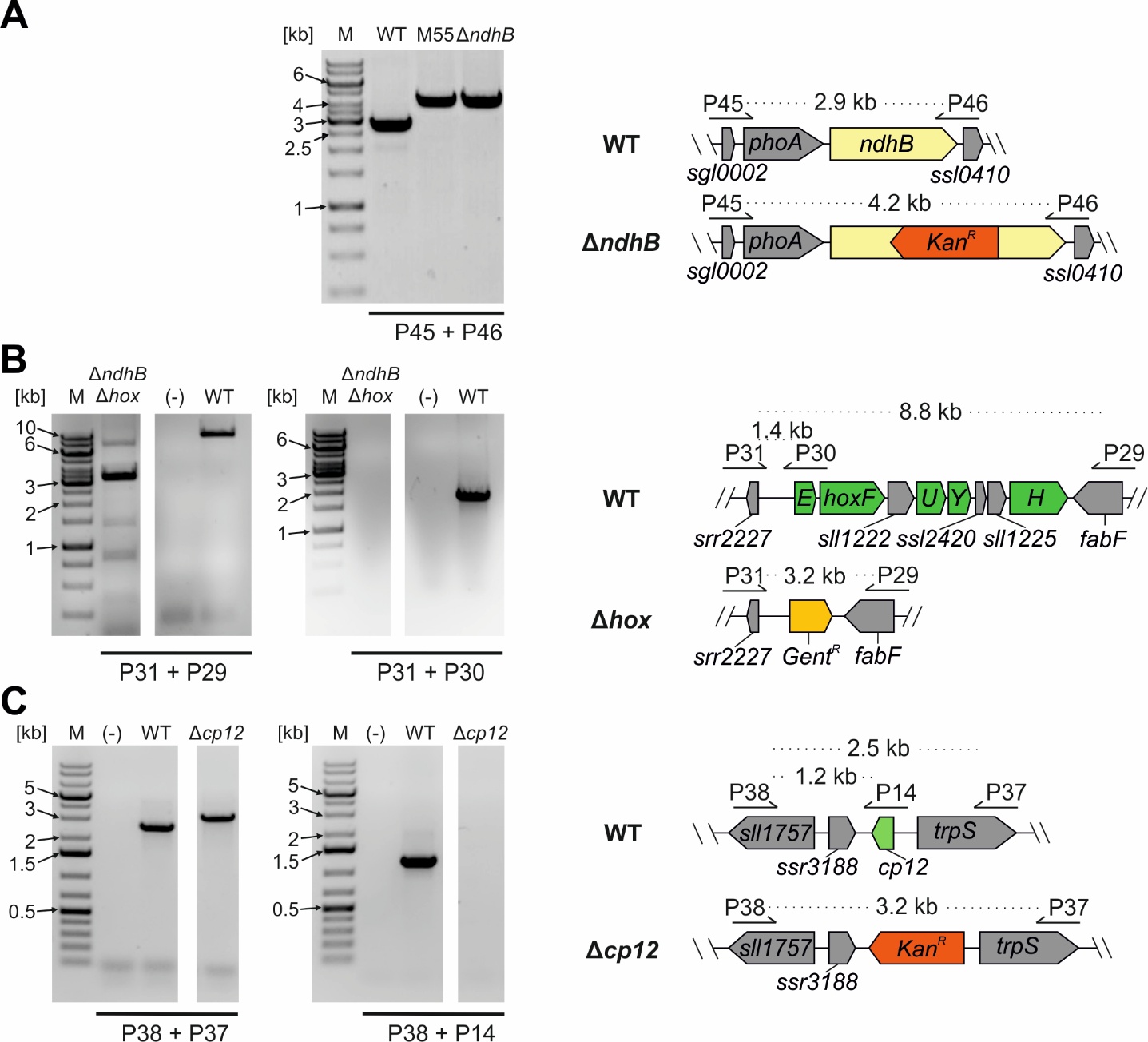

**Figure S1:** **Construction of deletion mutants of *Synechocystis*. A-C:** PCRs and schematic of the modified loci of the *ndhB* (A) interruption and *hox* (B) and *cp12* (C) deletion strains. Mutants were created by homologous recombination followed by selection on the respective antibiotic. The creation of the *ndhB* deletion strain is described in more detail elsewhere ^1^. Segregation of the strains was analyzed via PCR.

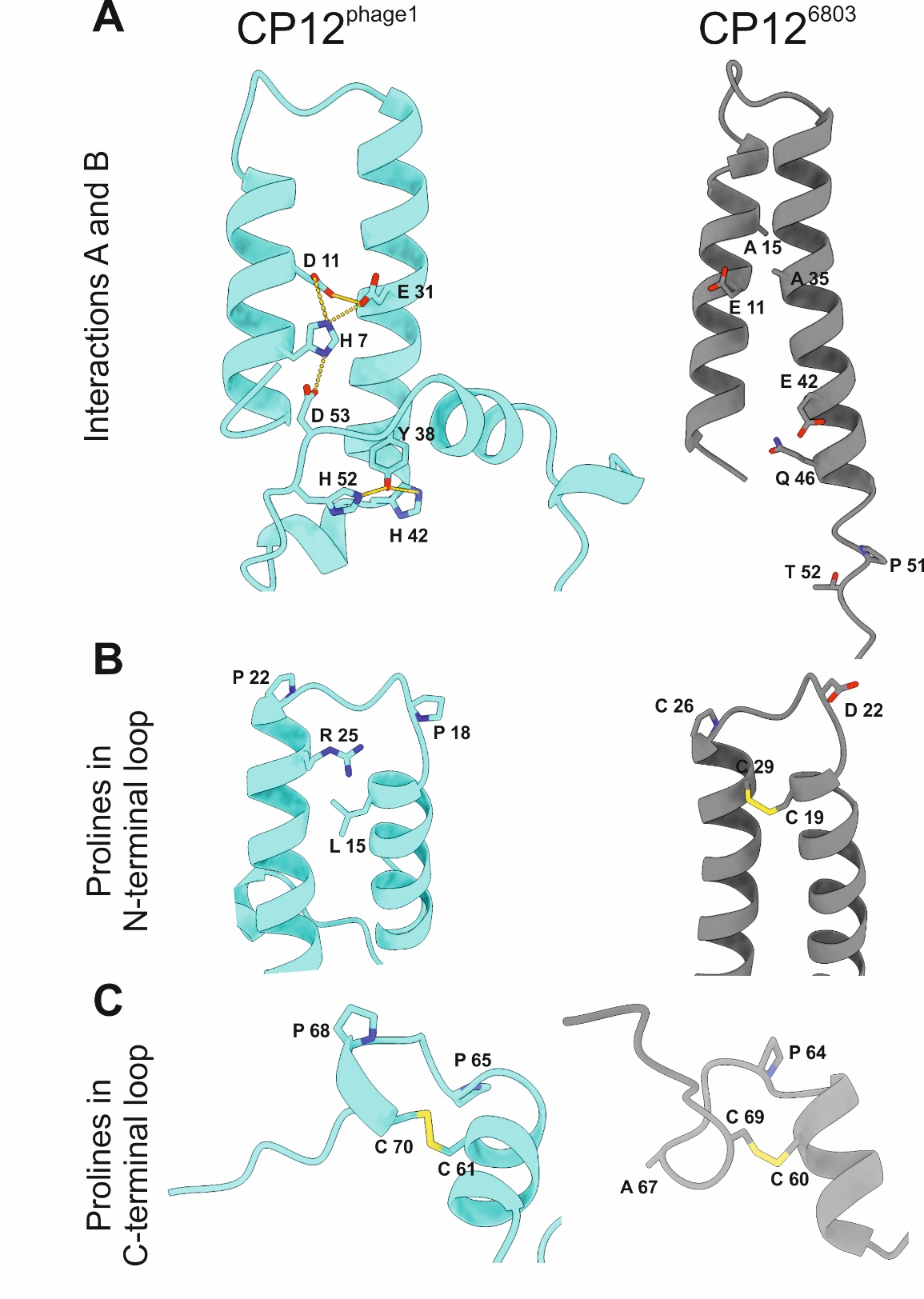

**Figure S2:** **Comparison of AlphaFold 3 structure predictions of CP12^phage1^ and CP12^6803^. A:** Predicted interaction network between residues unique to and conserved in phage-derived CP12 sequences and homologous positions in CP12^6803^ **B-C:** N-terminal (B) and C-terminal (C) loop in CP12^phage1^ and CP12^6803^**.** Proline and cysteine residues or respective positions in an alignment are presented with their sidechains.

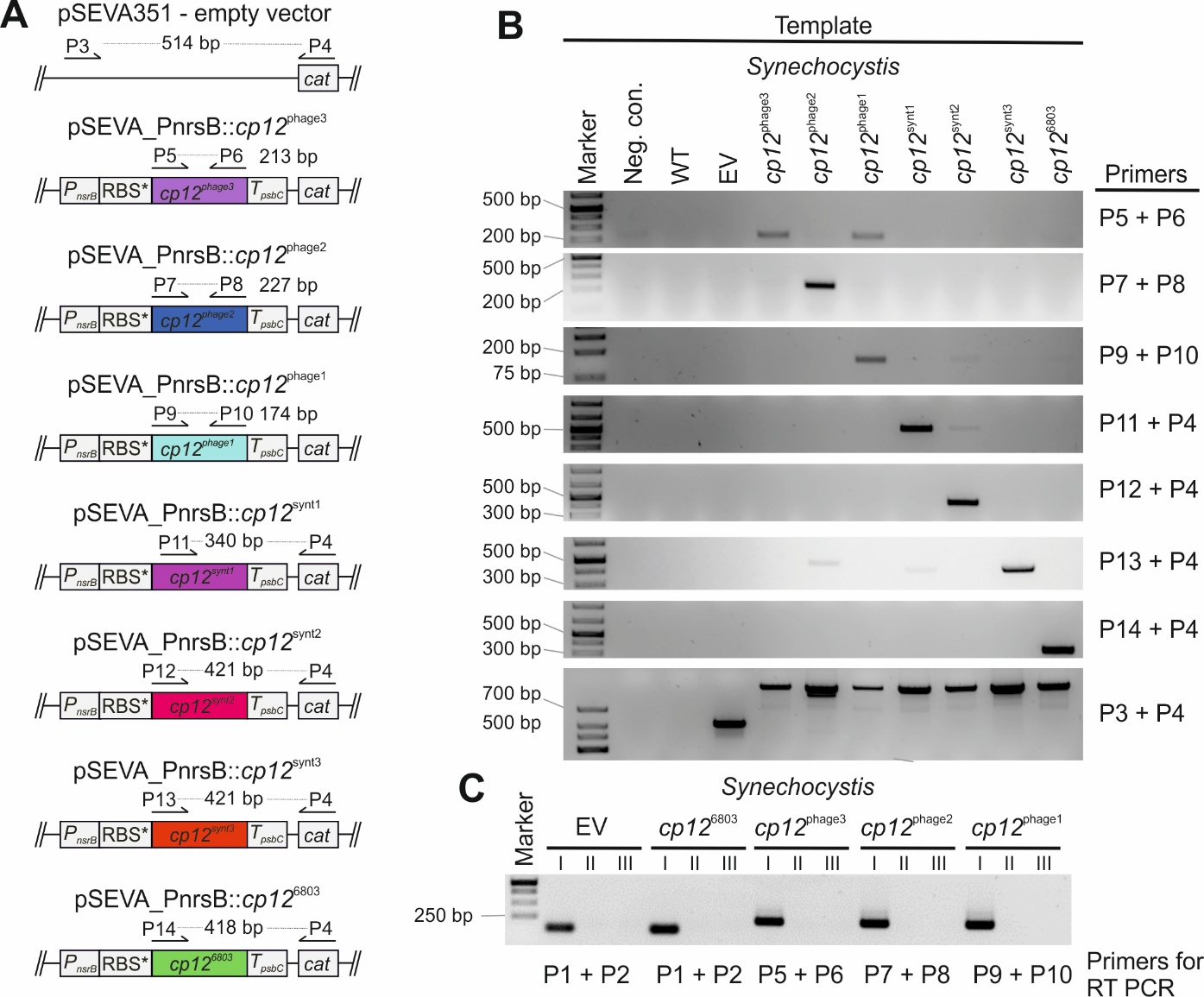

**Figure S3:** **Construction and verification of *ricp12* expressing mutants of *Synechocystis*. A:** Genetic configuration of the plasmids used for inducible expression of *cp12* variants. **B**: PCR confirming the insertion of the respective plasmids into *Synechocystis* WT. Additional bands for other strains are attributable to the only small differences between the sequences of some *cp12* variants. **C:** RT-PCR using cDNA as template (lanes I) confirming the transcription of representative *cp12* variants. For EV and *cp12*^6803^ the native CP12-mRNA was targeted (see Table S2). Additionally, PCR was performed on extracted RNA to ensure that it was not contaminated with gDNA (lanes II) and with water instead of template cDNA/RNA added as negative controls (lanes III).

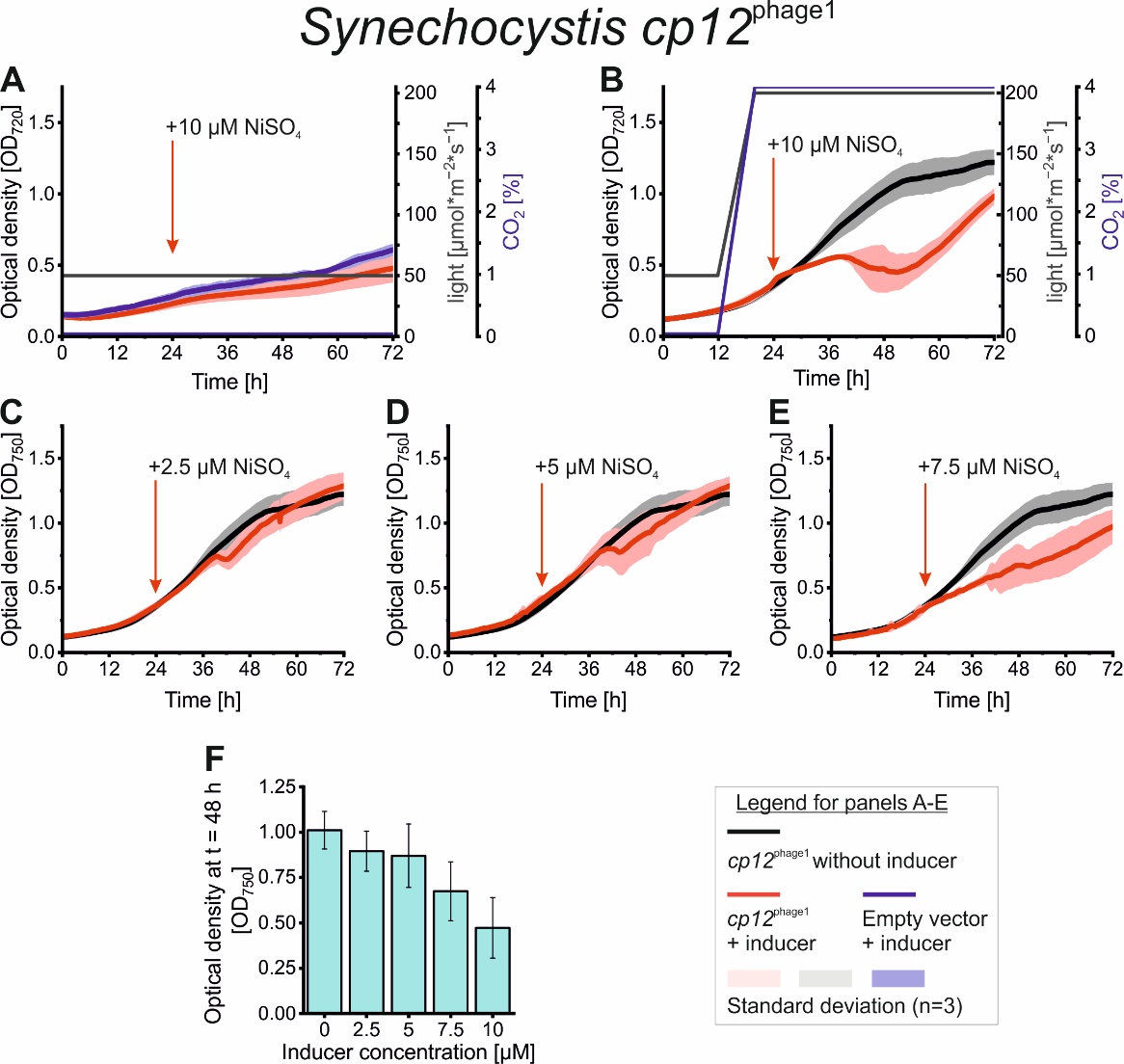

**Figure S4:** **Growth arrest upon induction of CP12^phage1^.** A-E: Growth of *Synechocystis* cp12^phage1^ in standard (A) or high light and high CO_2_ (B-E) conditions with different amounts of NiSO_4_ added as inducer of cp12^phage1^ expression at t = 24 h. Light intensity and CO_2_ concentration are depicted by black and blue lines, respectively. For high light and high CO_2_ conditions, both parameters were increased linearly from the standard conditions between 12 h and 20 h of the incubations. For direct comparison, growth curves with 10 µM inducer under high light and high CO_2_ conditions (B) from Fig. 5 are shown, again. Data show the smoothed average from automatic growth monitoring using the MC-1000 ± SD of three biological replicates F**:** Optical density of the cultures shown in B-E 24 h after induction of *cp12*^phage1^ with increasing amounts of the inducer (Ni^2+^). Data represent the average ± SD of three biological replicates.

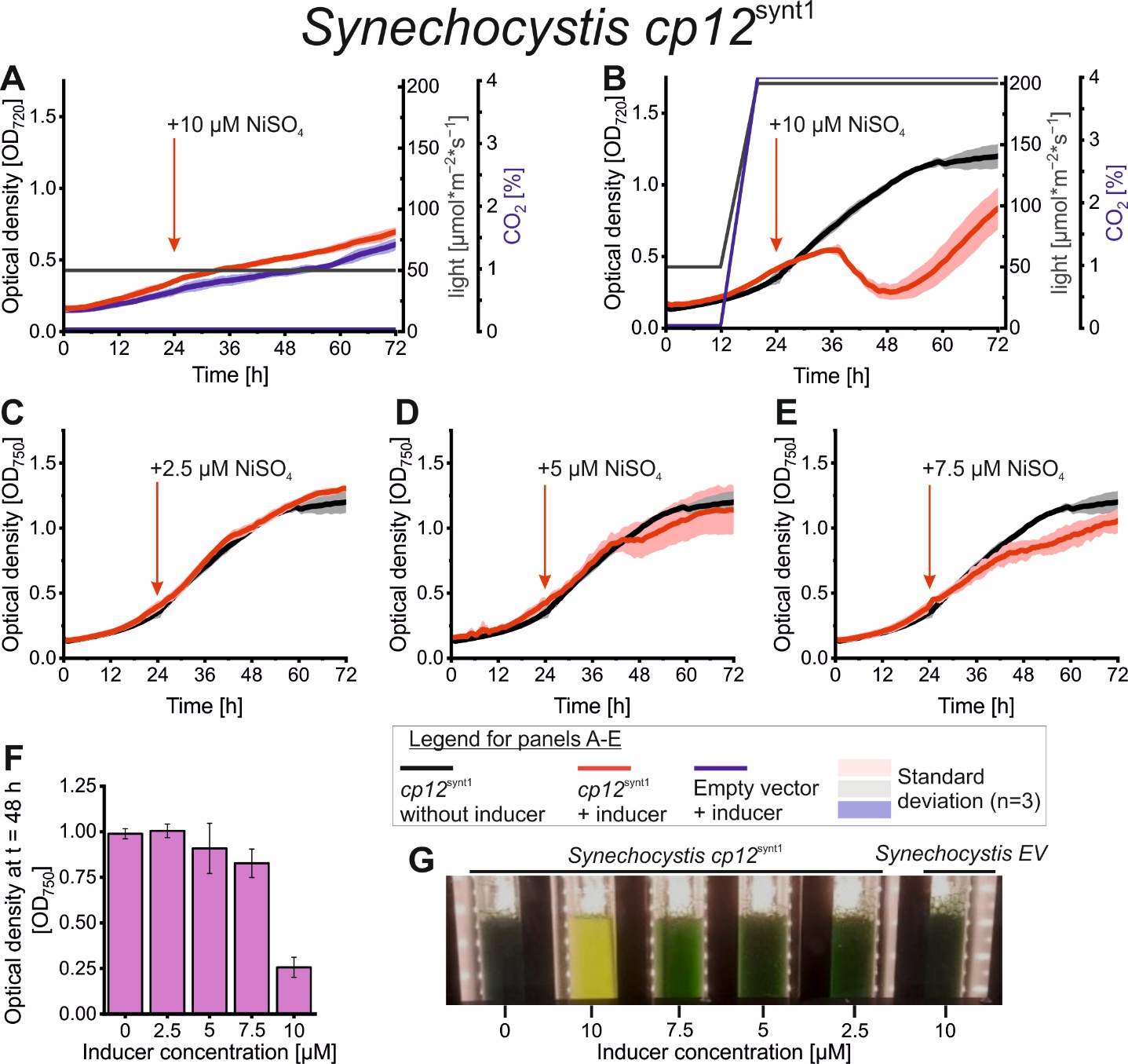

**Figure S5:** **Growth arrest upon induction of CP12^synt1^.** A-E: Growth of *Synechocystis* cp12^synt1^ in standard (A) or high light and high CO_2_ (B-E) conditions with different amounts of NiSO_4_ added as inducer of cp12^synt1^ expression at t = 24 h. For direct comparison, growth curves with 10 µM inducer under high light and high CO_2_ conditions (B) from Fig. 5 are shown, again. Light intensity and CO_2_ concentration are depicted by black and blue lines, respectively. For high light and high CO_2_ conditions, both parameters were increased linearly from the standard conditions between 12 h and 20 h of the incubations. Data show the smoothed average from automatic growth monitoring using the MC-1000 ± SD of three biological replicates F**:** Optical density of the cultures shown in B-E 24 h after induction of *cp12*^phage1^ with increasing amounts of the inducer (Ni^2+^). Data represent the average ± SD of three biological replicates. G: Photograph of cultures *Synechocystis* cp12^synt1^ and *Synechocystis* EV 24 h after addition of different amount of the inducer NiSO_4_.

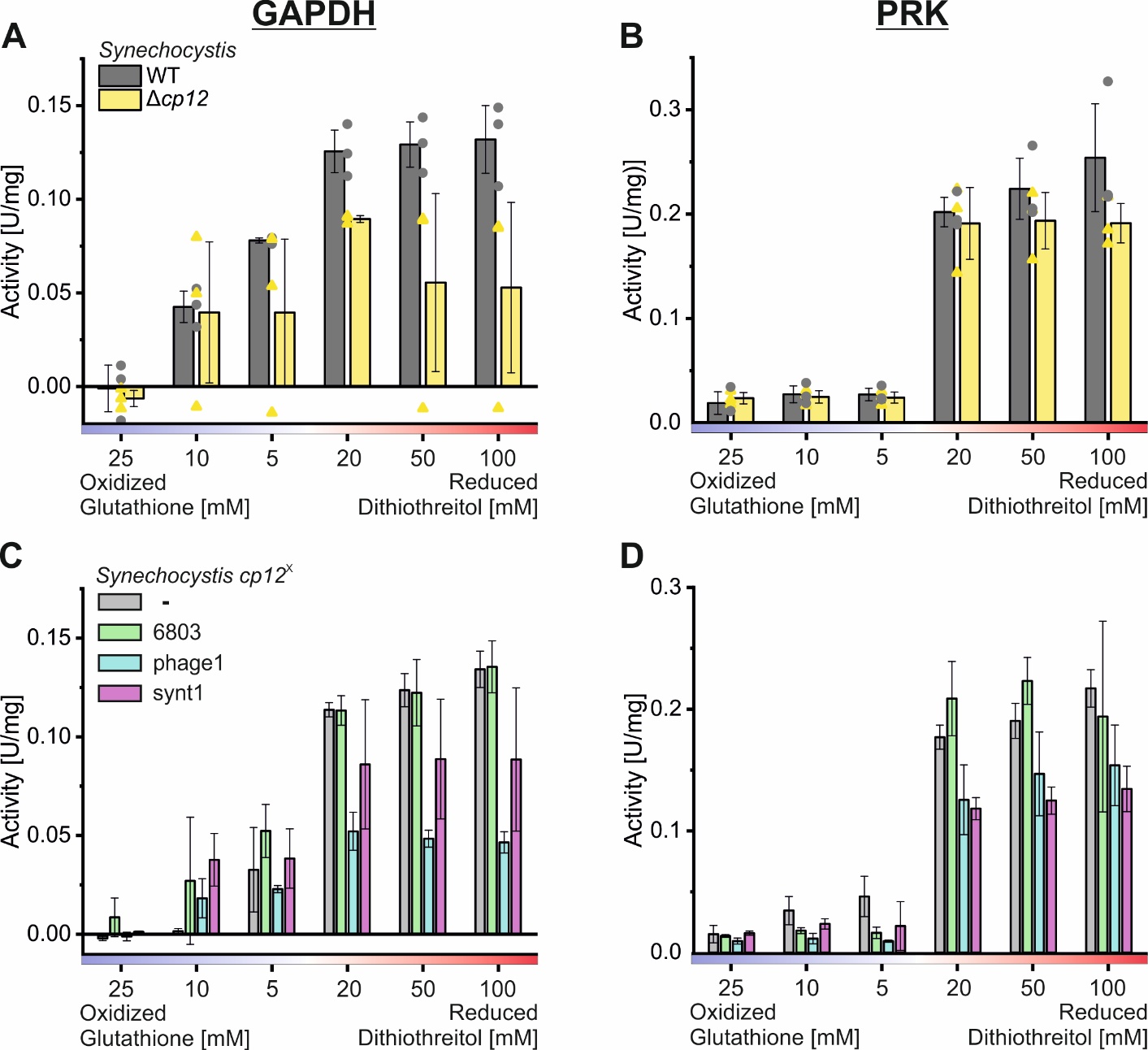

**Figure S6:** **GAPDH and PRK activity in different redox conditions A-B:** Enzymatic activity of glycerinaldehyde 3-phosphate dehydrogenase (GAPDH) (A) and phosphoribulokinase (PRK) (B) were analyzed in crude extracts of the *Synechocystis* WT or the *cp12* deletion mutant (Δ*cp12).* **C-D**: Enzymatic activity of GAPDH (C) and PRK (D) in crude extracts of *Synechocystis* strains harboring an empty vector (-) or expressing *cp12* variants 24 h after induction. Different redox states of the cell were recreated by the addition of different concentrations of oxidized glutathione or reduced dithiothreitol. Data represent average ± SD of three biological replicates. Activity: 1 Unit corresponds to an enzymatic turnover of 1 µmol min^-1^.

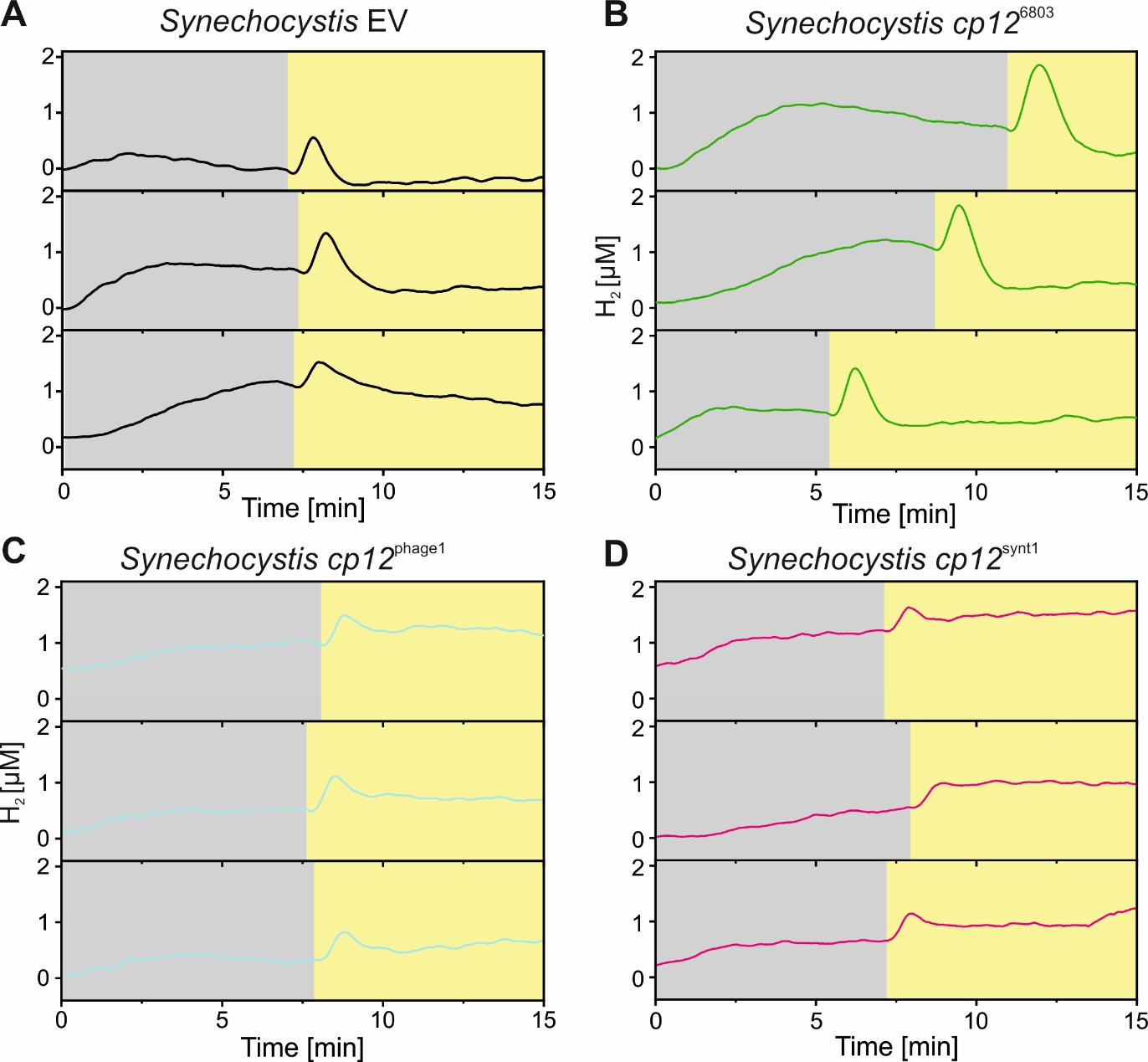

**Figure S7:** **H_2_ kinetics of *Synechocystis* strains expressing *ricp12* in a dark-light transition. A-D:** *Synechocystis* strains harboring an empty vector (EV) (A) or plasmids for the Ni^2+^-inducible expression of *cp12*^6803^ (B), *cp12*^phage1^ (C), or *cp12*^synth1^ (D) were cultivated photoautotrophically in conditions of 200 µE*m^-2^*s^-1^ light and 4 % CO_2_ and 10 µM of NiSO_4_ was added 8 – 12 h prior to the H_2_ production assay to induce *ricp12* expression. Equal amounts of cells by OD_750_ were then used to measure the H_2_ evolution using UniSense electrodes. To maintain anaerobic conditions also under photosynthetic conditions (light phase) cultures were supplemented with 10 mM glucose and an enzyme mixture of 40 U glucose oxidase and 50 U catalase (God/Cat). In the assay, cells were adapted to the dark until the concentration of fermentative H_2_ stabilized (grey background) than they were exposed to 200 µE*m^-2^*s^-1^ light (yellow background).

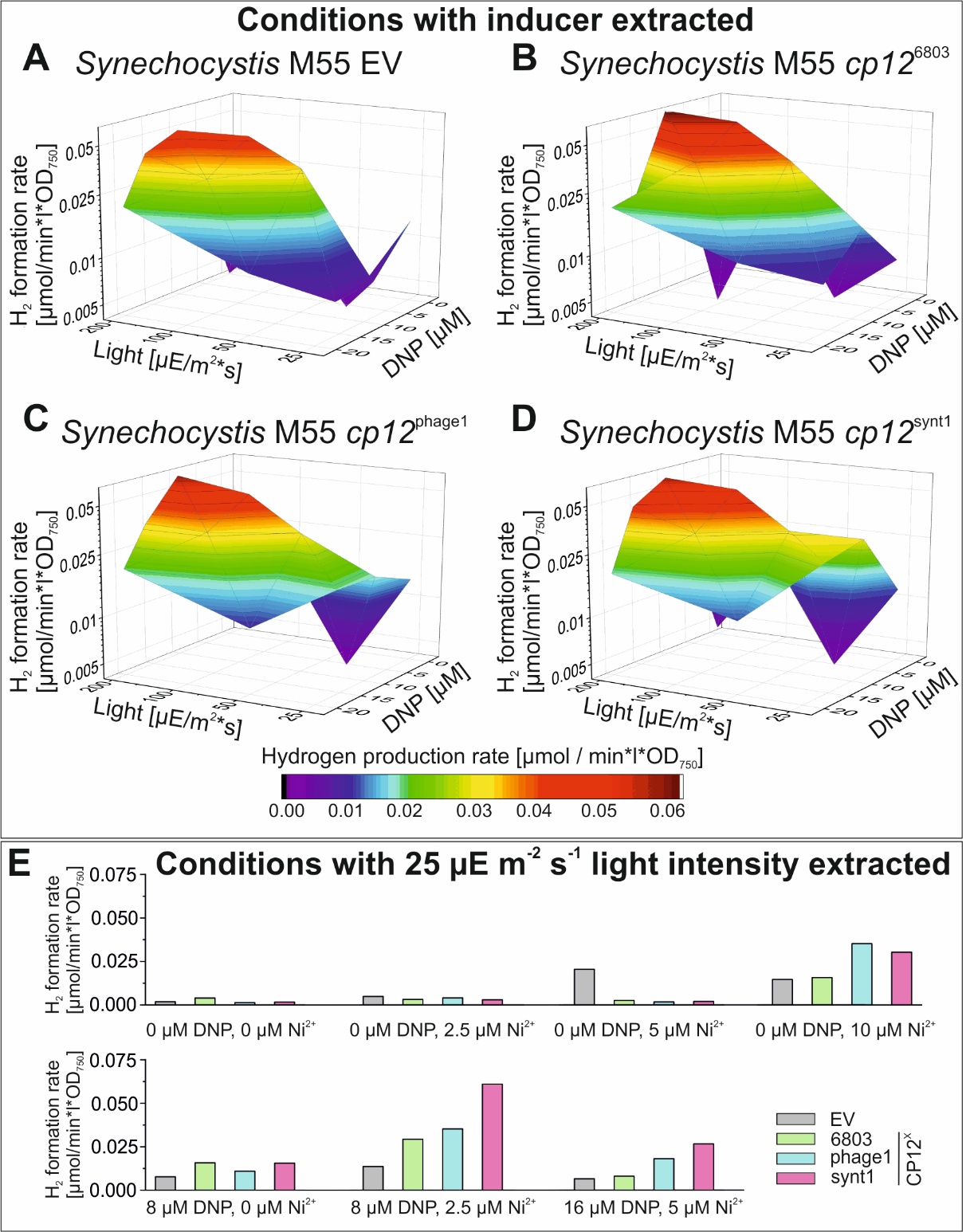

**Figure S8: Extracted results from a sparse matrix screening on the** **impact of light intensity, uncoupler concentration and riCP12 on H_2_ formation in the M55 mutant. A-D:** H_2_ formation rates of *Synechocystis* M55 strains harboring an empty vector (EV) (A) or plasmids for the Ni^2+^-inducible expression of *cp12*^6803^ (B), *cp12*^phage1^ (C), or *cp12*^synth1^ (D) calculated from GC-quantified H_2_ accumulating in the gas phase of sealed GC vials after 48 h of cultivation. Conditions with at least 2.5 µM Ni^2+^ (n = 25 per strain) added to the culture prior to transfer into the GC vials were extracted from the sparse matrix screening (Table S1). To maintain anaerobic conditions 10 mM glucose and an enzyme mixture of 40 U/ml glucose oxidase and 50 U/ml catalase (God/Cat) were added to the vials. **E:** H_2_ formation rates from the sparse matrix screening under low light conditions. Each set of bars represents one condition simultaneously tested once with all four strains. DNP: 2,4-dinitrophenol.

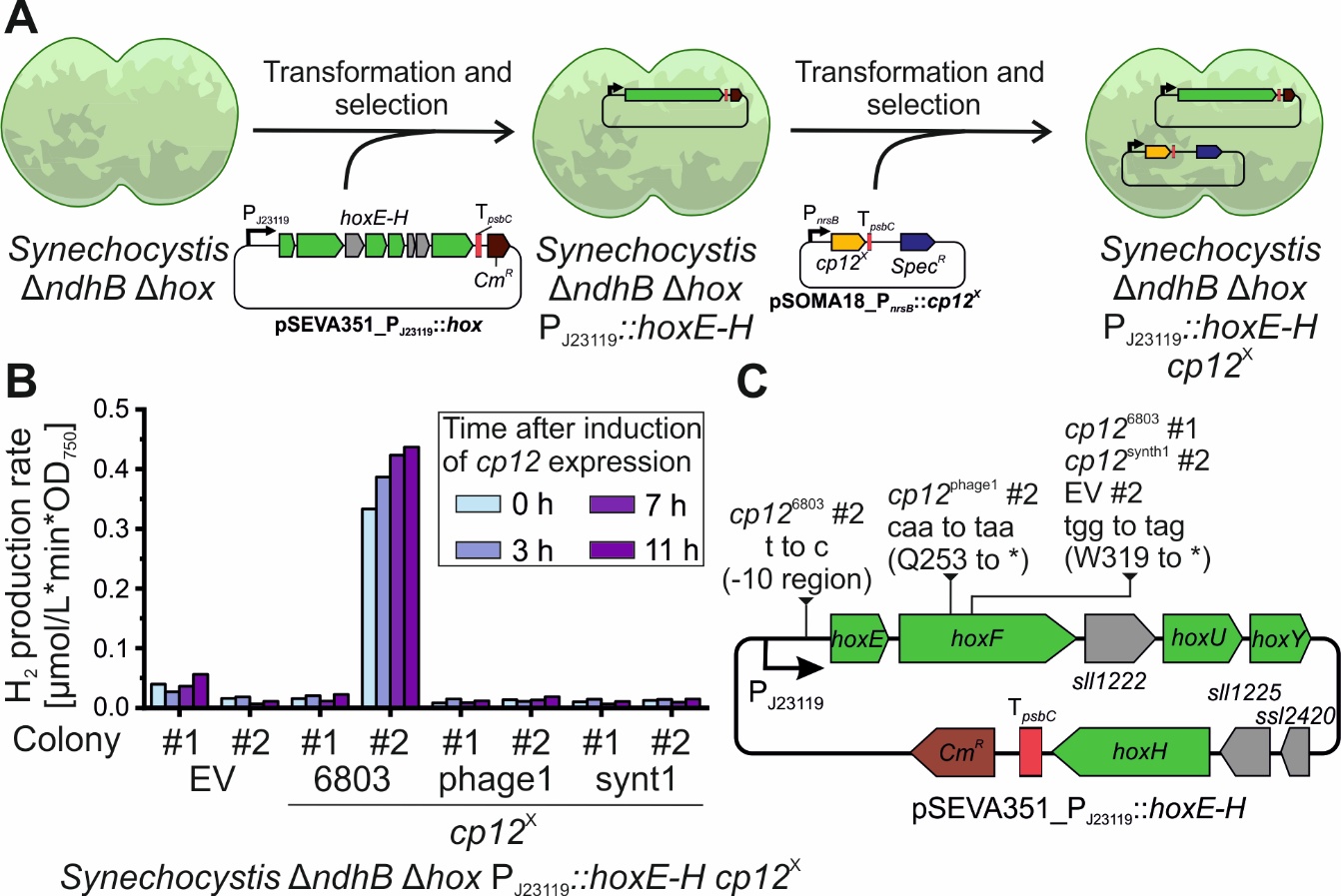

**Figure S9:** **Genetic instability of hox operon overexpressing *Synechocystis* Δ*ndhB* Δ*hox* strains. A:** Schematic of strain construction to produce *Synechocystis* Δ*ndhB* Δ*hox* P_J23119_::*hoxE-H* *cp12*^X^. **B:** H_2_ production rates calculated from GC-quantified H_2_ accumulating in the gas phase of sealed GC vials after ~24 h of cultivation. Prior to the start of the H_2_ production assay, cells were cultivated at conditions of 200 µE*m^-2^*s^-1^ light and 4 % CO_2_. Cells for H_2_ production quantification were taken from the cultures before and 3, 7, or 11 h after the addition of 5 µM of NiSO_4_. To maintain anaerobic conditions, GC vials were supplemented with *Pseudomonas* cells to an OD_600_ of 0.5 and 10 mM citrate. **C:** Mutations in plasmid for *hoxE-H* overexpression extracted from the *Synechocystis* Δ*ndhB* Δ*hox* P_J23119_::*hoxE-H* *cp12*^X^ cultures used for the H_2_ production assay. EV: empty vector control, t: thymine; a: adenine; g: guanine; c: cytosine; *: stop codon; t to c: mutation from T to C; caa to taa: mutation of CAA codon to TAA codon; tgg to tag: mutation of TGG codon to TAG codon.

**Table S1: Summary of H_2_ formation experiments using *Synechocystis*.** This table is provided as a separate CSV file that can be imported into standard spreadsheet software such as Microsoft Excel.

**Table S2: Sequence of primers used in this study.** The primer binding regions are given in bold, BpiI recognition sites are marked yellow, BsaI recognition sites are marked green, LguI recognition sites are marked cyan, recognition sites of Typ II restriction enzymes are marked red, cutting sites of the restriction enzymes are underlined, homologous regions for Gibson Assembly are marked in grey, and sequences of the promoter series J_23100_ are given in yellow letters. Oligos used to generate *in vitro* transcription templates of the RF00442 riboswitch constructs are partially overlapping to be used in an overlap-extension PCR (OE) to synthesize the corresponding full-length constructs.

| number | Primer name | Sequence 5´ -> 3´ | Primer target, Purpose |
| --- | --- | --- | --- |
| P1 | *cp12*^6803^ for | AAAGGTCTCGG**ATGAGCAATATTCAAGAAAAAATCG** | *cp12*^6803^ |
| P2 | *cp12*^6803^ rev | AAAGGTCTCGCTTT**CTAGTCGTCGTAAATGCGGC** | *cp12*^6803^ |
| P3 | pSEVA351 for | **CTAGCGCAGCGAATAGAC** | RSF1010 ori |
| P4 | pSEVA351 rev | **GGTGAAAGTTGGAACCTCTTACG** | P*cat* |
| P5 | *cp12*^phage3^ for | **TGGAATCCATTGAAAAACACATTG** | *cp12*^phage3^ |
| P6 | *cp12*^phage3^ rev | **CTAATCTTCGTACACCAAACATTCG** | *cp12*^phage3^ |
| P7 | *cp12*^phage2^ for | **ATGAAAACCATTGAAGAACACATTG** | *cp12*^phage2^ |
| P8 | *cp12*^phage2^ rev | **CTAATCATCGTACACCAAACATTCG** | *cp12*^phage2^ |
| P9 | *cp12*^phage1^ for | **GATGGAATCCATTGAACAACACA** | *cp12*^phage1^ |
| P10 | *cp12*^phage1^ rev | **TCCAAGGCGGTGGGATC** | *cp12*^phage1^ |
| P11 | *cp12*^synt1^ for | **TCAAGAAAAAATTGAACAAGTGTTGG** | *cp12*^synt1^ |
| P12 | *cp12*^synt2^ for | **CCGCCGAACGCCAAAAAC** | *cp12*^synt2^ |
| P13 | *cp12*^synt3^ for | **CCGCCTACCGCCAATAC** | *cp12*^synt3^ |
| P14 | *cp12*^6803^ for 2 | **ACCAACGTCAACAACATCCC** | *cp12*^6803^ |
| P15 | *cp12*^6803^ rev 2 | **CGTCGTAAATGCGGCACTC** | *cp12*^6803^ |
| P16 | PJ23100 for | AAAGGTCTCGGATGTTGACGGCTAGCTC**AGTCCT** | Produce PJ23100 |
| P17 | PJ23100 rev | AAAGGTCTCGCTTTGCTAGCACTGTACCT**AGGACT** | Produce PJ23100 |
| P18 | PJ23119 for | AAAGGTCTCGGATGTTGACAGCTAGCTC**AGTCCT** | Produce PJ23119 |
| P19 | PJ23119 rev | AAAGGTCTCGCTTTGCTAGCATTATACCT**AGGACT** | Produce PJ23119 |
| P20 | *hox* for 1 | AAAGGTCTCGG**ATGACCGTTGCCACCGATC** | *hoxE* |
| P21 | *hox* rev 1 | AAAGGTCTCGCAAA**TCTTCTTGGGTAGCTTCCC** | *hoxF* |
| P22 | *hox* for 2 | AAAGGTCTCGT**TTGATCAAACTAGAAAATCTTTG** | *hoxF* |
| P23 | *hox* rev 2 | AAAGGTCTCGCCC**TTGAATGATCGTTACCAAC** | *sll1222* |
| P24 | *hox* for 3 | AAAGGTCTCGAGG**GACCATTACCTAATTAGTAAC** | *sll1222* |
| P25 | *hox* rev 3 | AAAGGTCTCGCTTT**TTAATCCCGCTGGATGGAC** | *hoxH* |
| P26 | 5’ hom_*hox* for | AAAGCTCTTCAGTCGAAGACAAGCCA**CACCATTGCCACAGTAGAG** | *ssr2227* |
| P27 | 5’ hom_*hox* rev | AAAGCTCTTCAGGTGAAGACAACATC**CTGATTTTCACCCTTAGTCG** | Upstream of *hoxE* |
| P28 | 3’ hom_*hox* for | AAAGCTCTTCAGTCGAAGACAAAAAG**ACAAAAAACATTCAGACGGTC** | Downstream of *hoxH* |
| P29 | 3’ hom_*hox* rev | AAAGCTCTTCAGGTGAAGACAAGATG**TTTAAGCTTGTGGATTGGTG** | *fabF* |
| P30 | *hoxE* internal rev | **AAAAATTTCCTGGGCTTTATGCAG** | *hoxE* |
| P31 | Δ*hox* verification for | **CAAATTCCTTAGATGAACGC** | Upstream of *ssr2227* |
| P32 | P*nrsB* for | AAAGGTCTCGGATGT**TCCACCAGCAAAATTCGCA** | P*nrsB* |
| P33 | T*psbC* rev | AAAGGTCTCGCTTT**AACACCAGCGGGGAAAGG** | T*psbC* |
| P34 | 5’ hom_*cp12* for | AAAGCTCTTCAGTCGGTCTCGGCCA**ACGCCTCTGGATTTTGGG** | *sll1757* |
| P35 | 5’ hom_*cp12* rev | AAAGCTCTTCAGGTCTCGTAAC**CGCCAGGATTTCCAAAACG** | Upstream of *cp12* |
| P36 | 3’ hom_*cp12* for | AAAGCTCTTCAGTCGGTCTCGAAAG**AGAAGGAACCACAACGGG** | Downstream of *cp12* |
| P37 | 3’ hom_*cp12* rev | AAAGCTCTTCAGGTCTCGGATG**GCTTGAATTGGTTCCAAGGC** | *trpS* |
| P38 | Δ*cp12* verification for | **CTTGACCGGAGGCCAAAAG** | *sll1757* |
| P39 | Level 2 GGC for | AAAGCTCTTCATTT**CCGCCATGAGACCACGC** | GGC level 2 5’ insertion site |
| P40 | pSOMA17 bb rev | AAAGCTCTTCA**CGTTTTGCGCCTGTAGTGC** | pSC101 ori |
| P41 | SpecR for | AAAGCTCTTCAACG**GATGTCGCGCAGGCTGG** | Spec^R^ |
| P42 | SpecR rev | AAAGCTCTTCAAA**ACTAGATTTTAATGCGGATGTTGC** | Spec^R^ |
| P43 | M55 *ndhB* for | AAAGGTCTCGGCCA**GCAATTTGTCATTGGCACCGG** | *phoA* |
| P44 | M55 *ndhB* rev | AAAGGTCTCGGATG**GGGAGAACCTTCATAAACGTC** | *ndhB* |
| P45 | Δ*ndhB* verification for | **CTCTACCTCACCACCCTG** | *sgl0002* |
| P46 | Δ*ndhB* verification rev | **CATTGGCTTTTGCCTGCAAAGC** | *ssl0410* |

**Table S3: List of plasmids used and generated in this study.**

Plasmids were generated using Golden Gate Assembly (GGA) in which the given elements were fused via their BsaI, BpiI, or LguI cutting sites. Elements were added to the reactions as plasmids, synthesized linear dsDNA, or PCR products. PCR products are referred to as ‘PCR’ with the mentioning of the used primer pair as Pxx/Pxx and the used DNA template in brackets. If no template is mentioned, a PCR without a template yielding primer dimers resembling the desired sequence was carried out. Coding sequences (cds) of the *cp12* genes were synthesized by Eurofins genomics and are referred to as ‘ds *cp12*^X^’. Their full sequences are given in **Table S4**.

| **Vector** | **Level** | **5' overhang** | **3' overhang** | **Backbone** | **Selection** | **Plasmids or amplicons used in GGA to create the plasmid** | **Characteristics** | **Reference / source** |
| --- | --- | --- | --- | --- | --- | --- | --- | --- |
| **pGGC0** | 0 | GATG | AAAG | pUC18 | Amp^R^ |  | Level 0 empty entry vector based on pUC18 vector with additional integrated *BpiI/BsaI* restriction sites flanking *lacZα* | ^2^ |
| **pGGC1** | 1 | GCCA | GTTA | pUK21 | Kan^R^ |  | Level 1 position 1; empty entry vectors based on pUK21 vector with additional integrated BsaI/BpiI restriction sites flanking lacZα |  |
| **pGGC2.3** |  | GTTA | CTAG | pUK21 | Kan^R^ |  | Level 1 position 2.3; empty entry vectors based on pUK21 vector with additional integrated BsaI/BpiI restriction sites flanking *lacZα* and RBS* |  |
| **pGGC3** | 1 | CTAG | CAGA | pUK21 | Kan^R^ |  | Level 1 position 3; empty entry vectors based on pUK21 vector with additional integrated BsaI/BpiI restriction sites flanking *lacZα* |  |
| **pGGC11** | 0 | GATG | AAAG | pGGC0 | Amp^R^; Spec^R^ |  | Level 0 Spec^R^ |  |
| **pGGC17** | 1 | GTTA | CTAG | pGGC2.3 | Kan^R^ |  | Level 1 position 2.3 P*nrsB* with RBS* ^3^ |  |
| **pGGC21** | 1 | CAGA | TGTG | pGGC4 | Kan^R^ |  | Level 1 position 4 T*psbC* |  |
| **pGGC40** | 1 | GCCA | GTTA | pUK21 | Kan^R^ |  | End-linker level 1 spanning position 1 till 2 based on pUK21 vector with additional integrated *BsaI/BpiI* restriction sites flanking end-linker sequence TCGGTCACATGTGCATCCTCGATCTCA |  |
| **pGGC41** | 1 | CTAG | CATC | pUK21 | Kan^R^ |  | End-linker level 1 spanning position 3 till 7 based on pUK21 vector with additional integrated *BsaI/BpiI* restriction sites flanking end-linker sequence TCGGTCACATGTGCATCCTCGATCTCA |  |
| **pGGC43** | 1 | GAGC | CATC | pUK21 | Kan^R^ |  | End-linker level 1 spanning position 5 till 7 based on pUK21 vector with additional integrated *BsaI/BpiI* restriction sites flanking end-linker sequence TCGGTCACATGTGCATCCTCGATCTCA |  |
| **pGGC46** | 2 | GCCA | CATC | pBluescript II SK (+) | Amp^R^ |  | Level 2 empty entry vector based on pBluescript II SK (+) vector with additional integrated *BsaI* restriction sites flanking *lacZα* |  |
| **pGGC78** | P | GTTA | TGTG | pUK21 | Kan^R^ |  | Level P empty vector based on pUK21 vector with additional integrated BsaI/BpiI restriction sites flanking lacZα | ^4^ |
| **pGGC106** | 0 | GATG | AAAG | pUC18 | Amp^R^ | pGGC0; PCR P1/P2 (*Syn. 6803* gDNA) | Level 0 cds *cp12*^6803^ | This study |
| **pGGC107** | 1 | CTAG | CAGA | pUK21 | Kan^R^ | pGGC3; pGGC106 | Level 1 Position 3 cds *cp12*^6803^ | This study |
| **pGGC139** | 1 | GCCA | CTAG | pUK21 | Kan^R^ |  | End-linker level 1 spanning position 1 till 3 based on pUC19 vector with additional integrated *BsaI/BpiI* restriction sites flanking end-linker sequence TCGGTCACATGTGCATCCTCGATCTCA | ^4^ |
| **pGGC237** | 2 | - | - | pBluescript II SK (+) | Amp^R^, Kan^R^ | pGGC46; PCR P43/P44 (M55 gDNA) | Homologous recombination template for reconstruction of the *ndhB* interruption from the M55 mutant | ^1^ |
| **pGGC208** | 2 | GCCA | CATC | pSEVA351 | Cmc^R^ |  | Level 2 empty entry vector based on pSEVA351 vector with additional integrated *lacZα* flanked by *BsaI* restriction sites | ^2^ |
| **pGGC286** | 1 | GTTA | CTAG | pGGC2.3 | Kan^R^ |  | Level 1 Position 2 P_J23101_ with RBS* ^3^ | ^5^ |
| **pGGC317** | 2 | GCCA | CATC | pSOMA17 | Gent^R^ |  | Level 2 empty entry vector based on pSOMA17 vector with additional integrated *lacZα* flanked by *BsaI* restriction sites | ^2^ |
| **pGGC341** | 0 | GATG | AAAG | pGGC0 | Amp^R^, Gent^R^ |  | Level 0 Gent^R^::Tdouble | ^5^ |
| **pAI9** | 2 | - | - | pSEVA351 | Cmc^R^ | pGGC208; pGGC139; pGGC41 | pSEVA351 empty vector | This study |
| **pAI37** | 2 | - | - | pSEVA351 | Cmc^R^ | pGGC208; pGGC40; pGGC17; pAI49; pGGC21; pGGC43; | P*nrsB*:: *cp12*^phage3^::T*psbC* on pSEVA351 backbone | This study |
| **pAI38** | 2 | - | - | pSEVA351 | Cmc^R^ | pGGC208; pGGC40; pGGC17; pAI50; pGGC21; pGGC43; | P*nrsB*:: *cp12*^synt1^::T*psbC* on pSEVA351 backbone | This study |
| **pAI39** | 2 | - | - | pSEVA351 | Cmc^R^ | pGGC208; pGGC40; pGGC17; pAI51; pGGC21; pGGC43; | P*nrsB*:: *cp12*^phage1^::T*psbC* on pSEVA351 backbone | This study |
| **pAI40** | 2 | - | - | pSEVA351 | Cmc^R^ | pGGC208; pGGC40; pGGC17; pAI52; pGGC21; pGGC43; | P*nrsB*:: *cp12*^synt2^::T*psbC* on pSEVA351 backbone | This study |
| **pAI41** | 2 | - | - | pSEVA351 | Cmc^R^ | pGGC208; pGGC40; pGGC17; pAI53; pGGC21; pGGC43; | P*nrsB*:: *cp12*^synt3^::T*psbC* on pSEVA351 backbone | This study |
| **pAI42** | 2 | - | - | pSEVA351 | Cmc^R^ | pGGC208; pGGC40; pGGC17; pAI54; pGGC21; pGGC43; | P*nrsB*:: *cp12*^phage2^::T*psbC* on pSEVA351 backbone | This study |
| **pAI43** | 0 | GATG | AAAG | pUC18 | Amp^R^ | pGGC0; ds *cp12*^phage3^ | Level 0 cds *cp12*^phage3^ | This study |
| **pAI44** | 0 | GATG | AAAG | pUC18 | Amp^R^ | pGGC0; ds *cp12*^synt1^ | Level 0 cds *cp12*^synt1^ | This study |
| **pAI45** | 0 | GATG | AAAG | pUC18 | Amp^R^ | pGGC0; ds *cp12*^phage1^ | Level 0 cds *cp12*^phage1^ | This study |
| **pAI46** | 0 | GATG | AAAG | pUC18 | Amp^R^ | pGGC0; ds *cp12*^synt2^ | Level 0 cds *cp12*^synt2^ | This study |
| **pAI47** | 0 | GATG | AAAG | pUC18 | Amp^R^ | pGGC0; ds *cp12*^synt3^ | Level 0 cds *cp12*^synt3^ | This study |
| **pAI48** | 0 | GATG | AAAG | pUC18 | Amp^R^ | pGGC0; ds *cp12*^phage2^ | Level 0 cds *cp12*^phage2^ | This study |
| **pAI49** | 1 | CTAG | CAGA | pUK21 | Kan^R^ | pGGC3; pAI43 | Level 1 Position 3 cds *cp12*^phage3^ | This study |
| **pAI50** | 1 | CTAG | CAGA | pUK21 | Kan^R^ | pGGC3; pAI44 | Level 1 Position 3 cds *cp12*^synt1^ | This study |
| **pAI51** | 1 | CTAG | CAGA | pUK21 | Kan^R^ | pGGC3; pAI45 | Level 1 Position 3 cds *cp12*^phage1^ | This study |
| **pAI52** | 1 | CTAG | CAGA | pUK21 | Kan^R^ | pGGC3; pAI46 | Level 1 Position 3 cds *cp12*^synt2^ | This study |
| **pAI53** | 1 | CTAG | CAGA | pUK21 | Kan^R^ | pGGC3; pAI47 | Level 1 Position 3 cds *cp12*^synt3^ | This study |
| **pAI54** | 1 | CTAG | CAGA | pUK21 | Kan^R^ | pGGC3; pAI48 | Level 1 Position 3 cds *cp12*^phage2^ | This study |
| **pAI66** | 1 | GTTA | AAAG | pGGC2.0 | Kan |  | Level 1 position 2.0 Kan^R^::T*tonB* | ^5^ |
| **pAI135** | 2 | - | - | pSEVA351 | Cmc^R^ | pGGC208; pGGC40; pGGC17; pGGC107; pGGC21; pGGC43; | P*nrsB*:: *cp12*^6803^::T*psbC* on pSEVA351 backbone | This study |
| **pAI171** | - | GTC | ACC | pUC18 | Amp^R^ |  | pUC18 backbone with LguI sites flanking a *lacZα* | ^5^ |
| **pAI238** | 0 | GCCA | GATG | pUC18 | Amp^R^ | pAI171; PCR P26/P27 (*Syn. 6803* gDNA) | 5’ homologous region of *hox* operon | This study |
| **pAI239** | 0 | AAAG | CATC | pUC18 | Amp^R^ | pAI171; PCR P28/P29 (*Syn. 6803* gDNA) | 3’ homologous region of *hox* operon | This study |
| **pAI242** | P | GTTA | TGTG | pUK21 | Gent^R^, Kan^R^ | pGGC78; pAI238; pGGC341; pAI239 | Recombination template for deletion of the *hox* with Gent^R^::Tdouble resistance cassette | This study |
| **pAI244** | 0 | GATG | AAAG | pUC18 | Amp^R^ | pGGC0; PCR P16/P17 | Level 0 P_J23100_ | This study |
| **pAI245** | 0 | GATG | AAAG | pUC18 | Amp^R^ | pGGC0; PCR P18/P19 | Level 0 P_J23119_ | This study |
| **pAI246** | 0 | GATG | AAAG | pUC18 | Amp^R^ | pGGC0; PCR P20/P21 (*Syn. 6803* gDNA); PCR P22/P23 (*Syn. 6803* gDNA); PCR P24/P25 (*Syn. 6803* gDNA) | Level 0 cds *hoxE-H* | This study |
| **pAI247** | 1 | CTAG | CAGA | pUK21 | Kan^R^ | pGGC3; pAI246 | Level 1 Position 3 cds *hoxE-H* | This study |
| **pAI248** | 1 | GTTA | CTAG | pGGC2.3 | Kan^R^ | pGGC2.3; pAI244 | Level 1 Position 2 P_J23100_ with RBS* ^3^ | This study |
| **pAI250** | 1 | GTTA | CTAG | pGGC2.3 | Kan^R^ | pGGC2.3; pAI245 | Level 1 Position 2 P_J23119_ with RBS* ^3^ | This study |
| **pAI251** | 2 | - | - | pSEVA351 | Cmc^R^ | pGGC208; pGGC40; pGGC286; pAI247; pGGC21; pGGC43; | P_J23100_:: *hoxE-H*::T*psbC* on pSEVA351 backbone | This study |
| **pAI252** | 2 | - | - | pSEVA351 | Cmc^R^ | pGGC208; pGGC40; pAI248; pAI247; pGGC21; pGGC43; | P_J23101_:: *hoxE-H*::T*psbC* on pSEVA351 backbone | This study |
| **pAI253** | 2 | - | - | pSEVA351 | Cmc^R^ | pGGC208; pGGC40; pAI250; pAI247; pGGC21; pGGC43; | P_J23119_:: *hoxE-H*::T*psbC* on pSEVA351 backbone | This study |
| **pAI267** | 1 | GCCA | GTTA | pUC18 | Amp^R^ | pAI171; PCR P34/P35 (*Syn. 6803* gDNA) | 5’ homologous region of *cp12* | This study |
| **pAI268** | 1 | AAAG | CATC | pUC18 | Amp^R^ | pAI171; PCR P36/P37 (*Syn. 6803* gDNA) | 3’ homologous region of *cp12* | This study |
| **pAI269** | 2 | - | - | pBluescript II SK (+) | Kan^R^, Amp^R^ | pGGC46; pAI267, pAI66; pAI268 | Recombination template for *cp12* deletion with Kan^R^::Tdouble resistance cassette | This study |
| **pAI272** | 0 | GATG | AAAG | pUC18 | Amp^R^ | pGGC0; PCR P32/P33 (pAI38) | Level 0 P*nrsB*:: *cp12*^synt1^::T*psbC* | This study |
| **pAI273** | 0 | GATG | AAAG | pUC18 | Amp^R^ | pGGC0; PCR P32/P33 (pAI39) | Level 0 P*nrsB*:: *cp12*^phage1^::T*psbC* | This study |
| **pAI274** | 0 | GATG | AAAG | pUC18 | Amp^R^ | pGGC0; PCR P32/P33 (pAI135) | Level 0 P*nrsB*:: *cp12*^6803^::T*psbC* | This study |
| **pAI275** | 1 | GCCA | GTTA | pUK21 | Kan^R^ | pGGC1; pAI272 | Level 1 Position 1 P*nrsB*:: *cp12*^synt1^::T*psbC* | This study |
| **pAI276** | 1 | GCCA | GTTA | pUK21 | Kan^R^ | pGGC1; pAI273 | Level 1 Position 1 P*nrsB*:: *cp12*^phage1^::T*psbC* | This study |
| **pAI277** | 1 | GCCA | GTTA | pUK21 | Kan^R^ | pGGC1; pAI274 | Level 1 Position 1 P*nrsB*:: *cp12*^6803^::T*psbC* | This study |
| **pAI284** | 2 | - | - | pSEVA351 | Cmc^R^ | pGGC208; pAI275; pAI250; pAI247; pGGC21; pGGC43; | P*nrsB*:: *cp12*^synt1^::T*psbC_*P_J23119_:: *hoxE-H*::T*psbC* on pSEVA351 backbone | This study |
| **pAI285** | 2 | - | - | pSEVA351 | Cmc^R^ | pGGC208; pAI276; pAI250; pAI247; pGGC21; pGGC43; | P*nrsB*:: *cp12*^phage1^::T*psbC_*P_J23119_:: *hoxE-H*::T*psbC* on pSEVA351 backbone | This study |
| **pAI286** | 2 | - | - | pSEVA351 | Cmc^R^ | pGGC208; pAI277; pAI250; pAI247; pGGC21; pGGC43; | P*nrsB*:: *cp12*^6803^::T*psbC_*P_J23119_:: *hoxE-H*::T*psbC* on pSEVA351 backbone | This study |
| **pAI298** | 2 | GCCA | CATC | pSOMA18 | Spec^R^ | PCR P39/P40 (pGGC317); PCR P41/P42 (pGGC11) | Level 2 empty entry vector based on pSOMA18 vector with additional integrated *lacZα* flanked by *BsaI* restriction sites | This study |
| **pAI299** | 2 | - | - | pSOMA18 | Spec^R^ | pAI298; pGGC139; pGGC41 | pSOMA18 empty vector | This study |
| **pAI300** | 2 | - | - | pSOMA18 | Spec^R^ | pAI298; pGGC40; pGGC17; pGGC107; pGGC21; pGGC43; | P*nrsB*:: *cp12*^6803^::T*psbC* on pSOMA18 backbone | This study |
| **pAI301** | 2 | - | - | pSOMA18 | Spec^R^ | pAI298; pGGC40; pGGC17; pAI51; pGGC21; pGGC43; | P*nrsB*:: *cp12*^phage1^::T*psbC* on pSOMA18 backbone | This study |
| **pAI302** | 2 | - | - | pSOMA18 | Spec^R^ | pAI298; pGGC40; pGGC17; pAI50; pGGC21; pGGC43; | P*nrsB*:: *cp12*^synt1^::T*psbC* on pSOMA18 backbone | This study |

**Table S4: Synthetic gene sequences used in this study.** The given sequences were codon optimized for *Synechocystis* sp. PCC 6803 using JCat ^6^ and ordered as double stranded linear fragments from Eurofins genomics. BsaI recognition sites are marked green, cutting sites of the restriction enzymes are underlined, and the coding sequences are given in bold.

| Gene name | Sequence |
| --- | --- |
| *cp12*^phage1^ | CAGTAGGTCTCGG**ATGGAATCCATTGAACAACACATTGAAAAAGATAAAGAAATTTTGGATAACCCCACCACCTCCCCCCAAGCCCGCCGCCACATTGAAGGCGAATTGCACGAATTGGAAGAATACGTGGAACACCACAAAAAAGAAATTGAAGCCGGCGATCACCACGATCCCACCGCCTTGGAATTGTTTTGTGATCAAAACCCCTCCGAACCCGAATGTTTGGTGTACGATGATTAG**AAAGCGAGACCATGCT |
| *cp12*^phage2^ | CAGTAGGTCTCGG**ATGAAAACCATTGAAGAACACATTGAAAAAGATCACGCCATTTTGGATGATCCCACCACCTCCCCCGCCGCCCGCCGCCACTACAAAGAAGAATTGCACGAATTGGAAGTGTACCACAACCACCACCCCGAAGATCACCACGATCCCAACGCGCTCGAATTGTTCTGCGAAATGCACCCCGATGAACCCGAATGTTTGGTGTACGATGATTAG**AAAGCGAGACCATGCT |
| *cp12*^phage3^ | CAGTAGGTCTCGG**ATGGAATCCATTGAAAAACACATTGAAAAAGATAAAGAAATTTTGGATAACCCCATGATTTCCCCCAACCAACGCCGCCACATTGAAGGCGAATTGCACGAATTGGAAGATTACGCCGAACACCACAAAAAAGAAATTGAAGCCGGCGATCACCACGATCCCTCCCCCTTGGAATTGTACTGTGATGCCAACCCCTCCGAACCCGAATGTTTGGTGTACGAAGATTAG**AAAGCGAGACCATGCT |
| *cp12*^synt1^ | CAGTAGGTCTCGG**ATGTCCTTTTCCAACATTCAAGAAAAAATTGAACAAGTGTTGGCCAACGCCCGCCAAGTGTCCTCCACCGATGAAGCCTCCCCCGCCGAAGCCAAAGCCGTGTGGGATGAAGTGGAAGAATTGGAAGCCGAATTTGAAGCCGCCCGCCAATACCACCCCACCCAAACCACCCACGAAAAATTTGATGATGAAAACCCCGATTCCGAAGAATCCCGCGATTACGATGATTAG**AAAGCGAGACCATGCT |
| *cp12*^synt2^ | CAGTAGGTCTCGG**ATGGCCGAATCCCGCATTCAAGAAAAAATTGAACAAGAAAAAGCCAACGCCCGCCAACGCTCCTCCACCGATGAAGCCTCCCCCGCCGAAGCCGCCGCCGCCTGGGATGCCGTGGAAGAATTGGAAGCCGAATTGGCCGCCGAACGCCAAAAACACCCCACCCAAACCACCGAAGAAAAATTTGATGATGAAAACCCCGATTACGCCGAAAAACGCGATTACGATGATTAG**AAAGCGAGACCATGCT |
| *cp12*^synt3^ | CAGTAGGTCTCGG**ATGGCCTTTTCCCGCATTCAAGAAAAAATTGAACAAGAAAAAGCCAACGCCCGCCAACGCTCCTCCACCGATGAAGCCTCCCCCGCCGAAGCCGCCGCCGCCTGGGATGCCGTGGAAGAATTGGAAGCCGAATTGGCCGCCTACCGCCAATACCACCCCACCCAAACCACCGAAGAAAAATTTGATGATGAAAACCCCGATTACGCCGAAAAACGCGATTACGATGATTAG**AAAGCGAGACCATGCT |

**Table S5: List of strains used and generated in this study.**

| **Name** | **Chromosomal Adaptations** | **Replicative Plasmids** | **Background / Origin** |
| --- | --- | --- | --- |
| *Escherichia coli* TOP10 |  |  | ThermoFisher |
| *Pseudomonas taiwanensis* VLB120 ΔpSTY |  | Cured megaplasmid pSTY | ^7^ |
| *Synechocystis sp.* PCC 6803 - (WT) |  |  | Pasteuer Culture Collection of Cyanobacteria |
| *Synechocystis sp.* PCC 6803 M55 - (M55) | Δ*ndhB* |  | ^8^ |
| *Synechocystis sp.* PCC 6803 Δ*hoxH* | Δ*hoxH* |  | ^9^ |
| *Synechocystis sp.* PCC 6803 Δ*ndhB* | Δ*ndhB* |  | ^1^ |
| *Synechocystis sp.* PCC 6803 Δ*cp12* | Δ*cp12* |  | WT |
| *Synechocystis sp.* PCC 6803 Δ*ndhB* Δ*hox* | Δ*ndhB* Δ*hox* |  | *Synechocystis sp.* PCC 6803 Δ*ndhB* |
| *Synechocystis sp.* PCC 6803 EV |  | pAI9 | WT |
| *Synechocystis sp.* PCC 6803 *cp12*^6803^ |  | pAI135 | WT |
| *Synechocystis sp.* PCC 6803 *cp12*^phage1^ |  | pAI39 | WT |
| *Synechocystis sp.* PCC 6803 *cp12*^phage2^ |  | pAI42 | WT |
| *Synechocystis sp.* PCC 6803 *cp12*^phage3^ |  | pAI37 | WT |
| *Synechocystis sp.* PCC 6803 *cp12*^synt1^ |  | pAI38 | WT |
| *Synechocystis sp.* PCC 6803 *cp12*^synt2^ |  | pAI40 | WT |
| *Synechocystis sp.* PCC 6803 *cp12*^synt3^ |  | pAI41 | WT |
| *Synechocystis sp.* PCC 6803 M55 EV | Δ*ndhB* | pAI9 | M55 |
| *Synechocystis sp.* PCC 6803 M55 *cp12*^6803^ | Δ*ndhB* | pAI135 | M55 |
| *Synechocystis sp.* PCC 6803 M55 *cp12*^phage1^ | Δ*ndhB* | pAI39 | M55 |
| *Synechocystis sp.* PCC 6803 M55 *cp12*^synt1^ | Δ*ndhB* | pAI38 | M55 |
| *Synechocystis sp.* PCC 6803 Δ*ndhB* Δ*hox* P_J23100_::*hoxE-H* | Δ*ndhB* Δ*hox* | pAI251 | *Synechocystis sp.* PCC 6803 Δ*ndhB* Δ*hox* |
| *Synechocystis sp.* PCC 6803 Δ*ndhB* Δ*hox* P_J23101_::*hoxE-H* | Δ*ndhB* Δ*hox* | pAI252 | *Synechocystis sp.* PCC 6803 Δ*ndhB* Δ*hox* |
| *Synechocystis sp.* PCC 6803 Δ*ndhB* Δ*hox* P_J23119_::*hoxE-H* | Δ*ndhB* Δ*hox* | pAI253 | *Synechocystis sp.* PCC 6803 Δ*ndhB* Δ*hox* |
| *Synechocystis sp.* PCC 6803 Δ*ndhB* Δ*hox* P_J23119_::*hoxE-H*_*cp12*^6803^ | Δ*ndhB* Δ*hox* | pAI286 | *Synechocystis sp.* PCC 6803 Δ*ndhB* Δ*hox* |
| *Synechocystis sp.* PCC 6803 Δ*ndhB* Δ*hox* P_J23119_::*hoxE-H*_*cp12*^phage1^ | Δ*ndhB* Δ*hox* | pAI285 | *Synechocystis sp.* PCC 6803 Δ*ndhB* Δ*hox* |
| *Synechocystis sp.* PCC 6803 Δ*ndhB* Δ*hox* P_J23119_::*hoxE-H*_*cp12*^synth1^ | Δ*ndhB* Δ*hox* | pAI284 | *Synechocystis sp.* PCC 6803 Δ*ndhB* Δ*hox* |
| *Synechocystis sp.* PCC 6803 Δ*ndhB* Δ*hox* P_J23119_::*hoxE-H* EV | Δ*ndhB* Δ*hox* | pAI253 pAI299 | *Synechocystis sp.* PCC 6803 Δ*ndhB* Δ*hox* P_J23119_::*hoxE-H* |
| *Synechocystis sp.* PCC 6803 Δ*ndhB* Δ*hox* P_J23119_::*hoxE-H cp12*^6803^ | Δ*ndhB* Δ*hox* | pAI253 pAI300 | *Synechocystis sp.* PCC 6803 Δ*ndhB* Δ*hox* P_J23119_::*hoxE-H* |
| *Synechocystis sp.* PCC 6803 Δ*ndhB* Δ*hox* P_J23119_::*hoxE-H cp12*^phage1^ | Δ*ndhB* Δ*hox* | pAI253 pAI301 | *Synechocystis sp.* PCC 6803 Δ*ndhB* Δ*hox* P_J23119_::*hoxE-H* |
| *Synechocystis sp.* PCC 6803 Δ*ndhB* Δ*hox* P_J23119_::*hoxE-H cp12*^synt1^ | Δ*ndhB* Δ*hox* | pAI253 pAI302 | *Synechocystis sp.* PCC 6803 Δ*ndhB* Δ*hox* P_J23119_::*hoxE-H* |
